# Variation in response to water availability across *Phlox* species

**DOI:** 10.1101/2025.06.17.660163

**Authors:** Christina Steinecke, Julius A. Tabin, James Caven, Charles O. Hale, Antonio Serrato-Capuchina, Robin Hopkins

**Affiliations:** Department of Organismic and Evolutionary Biology, Harvard University, Cambridge, Massachusetts 02138 USA; Arnold Arboretum of Harvard University, Roslindale, Massachusetts 02131 USA; Section of Plant Breeding and Genetics, Cornell University, Ithaca, New York 14853 USA; Department of Biology, Boston College, Boston, Massachusetts 02467

**Keywords:** Phenotypic plasticity, *Phlox*, niche modeling, ecological differentiation, life history

## Abstract

Plants adapt to environmental variation both by evolving divergent trait means and by plastically adjusting trait expression in response to local conditions. While these dual strategies are essential for persistence in diverse environments, how they interact and vary across closely related species is understudied. For plants, water availability is a particularly important selective force that shapes species distributions, selects for growth habit and life history strategy, and can dictate individuals’ plastic expressions of trait values and reproductive success. Here, we use ecological niche modeling, field soil collections, and a controlled drought experiment to test how biogeography and evolutionary history influence responses to limited water availability in three closely related Texas annual *Phlox* species and their F1 hybrids. We infer that the species occupy distinct niches that diverge along a primary axis of water availably and soil moisture. Each species has a distinct vegetative growth habit that does not match broad predictions of divergence in response to water availability. Nevertheless, we find that all the species show a significant morphological response to controlled soil dry down with reduced biomass, smaller leaves, and fewer flowers, as would be predicted in a response to drought. We find that *Phlox drummondii*, which occupies intermediate habitats, exhibits the strongest plastic response to water limitation, despite it not having the broadest environmental niche. Additionally, most hybrids involving *P. drummondii* display intermediate phenotypes in both wet and dry treatments, while hybrids between *P. cuspidata* and *P. roemeriana* show phenotypes consistent with hybrid vigor. These results challenge the hypothesis that species from broader environments evolve greater plasticity. Instead, the most plastic species did not have the broadest niche, suggesting plasticity and niche breadth may evolve independently.

## INTRODUCTION

As sessile organisms, plants must adapt to and grow in the environment in which they germinate. This can be done by both evolving divergent traits that optimize performance in a particular environment, and through responding to the environment by adjusting traits for higher performance (Bohnert et al., 1995; Kawecki and Ebert, 2004; van Kleunen and Fischer, 2005; Anderson et al., 2011; Stotz et al., 2021; Vinton et al., 2022). These two mechanisms of divergence and phenotypic plasticity are widely understood to be important for organisms (particularly sessile plants) to survive and thrive in extreme and changing environments (Anderson et al., 2011; Stotz et al., 2021). Despite extensive work on each of these strategies, how they interact and contribute to environmental responses across clades of closely related species is still poorly understood, particularly in empirical systems (but see Ghalambor et al., 2007; Pfennig et al., 2010; Husemann et al., 2017).

Phenotypic plasticity broadly refers to any environmentally induced trait variation including physiological, developmental, morphological, or behavioral traits (West- Eberhard, 1989; DeWitt and Scheiner, 2004). Whereas divergence is heritable trait variation that differentiates organisms adapted to different environments, plasticity refers to the ability of an organism or genotype to produce multiple phenotypes depending on environmental conditions encountered (Bradshaw, 1965; Via et al., 1995; Roff, 1999; Agrawal 2001). In this way, plasticity can be a “first responder” that enables organisms to rapidly change and survive in new or fluctuating environments. Although widely studied in plants, plasticity is universal across living organisms and manifests at all levels of biological organization from gene expression to physiology, morphology, and even life history strategy.

The magnitude of a plastic response can vary in intensity or extent across individuals, populations, and species, even those that are closely related. In much the same way that trait means evolve and respond to selection from the environment to optimize fitness, the degree of plasticity can also evolve and therefore vary across organisms (Wund et al., 2008; Nielson and Papaj, 2022; Vinton et al., 2022). Given the ubiquity of phenotypic plasticity, it is important to understand how it varies. From an ecological perspective, it has been hypothesized that species which inhabit broad abiotic or biotic environmental conditions that show high levels of stochasticity will have greater plasticity then species that are restricted to specific environmental niches (Pfenning et al., 2010; Leung et al., 2020; Stotz et al., 2021).

The patterns of variation in trait plasticity and trait means across populations and species can reflect the potential for selective response to the environment (Fox et al., 2019; Lie et al., 2022). Plasticity is an initial response to environmental stimuli and, when selective conditions persist, initially plastic traits may become genetically assimilated and canalized (Agrawal et al., 1999; Chun et al., 2007; Wund et al., 2008). This model of evolution predicts patterns of how variation in trait means and trait plasticity are distributed across closely related species. Specifically, the dominant axis of genetic divergence in response to environmental variation is predicted to be parallel to the plastic response to environmental variation (Lind et al., 2015). In other words, the plastic response of a species to an environmental stimulus is hypothesized to be in the same direction as trait divergence in closely related species that are adapted across the gradient of that same environment (de Jong, 2005; Radersma et al., 2020). For instance, trait divergence is closely associated with distinct elevational changes in closely related species of *Senecio* (Walter et al., 2022). In general, ecology (Kulkarni et al., 2011), evolutionary history (Pigliucci et al., 1999; Kellermann et al., 2018), and environmental stochasticity (Leung et al., 2020) determine the extent of plastic responses in nature.

Much can be inferred about the evolution of plasticity and adaptive divergence by comparing closely related species; and even more can be learned by also examining hybrids. Hybridization between lineages that differ in adaptive trait means or in plastic responses to environmental conditions could result in hybrid individuals with either intermediate or extreme (transgressive) trait means or plasticity (Rieseberg et al., 1999; Stelkens and Seehausen, 2009; Dittrich-Reed and Fitzpatrick, 2012; Husemann et al., 2017). These two alternative outcomes could suggest different evolutionary paths.

Intermediate trait means in hybrids suggest species differences are due to additive genetic variation, consistent with evolution from shared ancestral variation that does not interact with strong negative or positive epistasis (Stelkens and Seehausen, 2009).

Similar inferences could be made about additive genetic variation underlying intermediate plasticity in hybrids. In hybrids, transgressive traits are hypothesized to emerge when parental species evolve novel genetic variation across multiple loci or pathways in isolation that either complement each other’s phenotypic effects or interact with epistasis in hybrids (Rieseberg et al., 1999; Rieseberg et al., 2003; Stelkens and Seehausen, 2009; Kagawa and Takimoto, 2017). In this way, investigations into the environmental responses of hybrid lineages can be a tractable way to gain insight into the evolution of environmental response.

Water availability is one of the most important abiotic environmental determinants of plant success. Variation in water availability across time and space significantly contributes to regulating species presence and absence and range limits (e.g., in Eckstein, 2005, van Kleunen and Fischer, 2005, and Liu et al., 2017). Furthermore, there are countless studies documenting the plastic response to drought across all scales of biological organization from gene expression to developmental timing, to morphology (Des Marais et al., 2013). There is extensive literature on the response of model and agricultural species to drought (El Hafid et al., 1998; Bartlett et al., 2016; Hoover et al., 2017), but we know much less about how drought response evolves in both non-model species and across closely related species. Plants have likely evolved numerous unique responses to mitigate drought stress, including genetic, physiological, and morphological adaptations. Typically, drought response manifests as alterations in overall growth, membrane integrity, photosynthetic activity, and water-use efficiency (Hoover et al., 2017; Gupta et al., 2020). Experimental comparative studies of species across moisture gradients thus have the potential to further tease apart how species adapt to drought stress and the underlying genetics of these drought response traits.

*Phlox drummondii*, *Phlox cuspidata*, and *Phlox roemeriana* comprise a small clade of three annual herbaceous flowering plant species native to Texas, USA. These three species are recently diverged (∼2 mya), with *P. cuspidata* being an outgroup to sibling taxa *P. drummondii* and *P. roemeriana* (Roda et al., 2017, Garner et al., 2024; Figure 1A). Among these, *P. drummondii* occupies a central range, overlapping with *P. cuspidata* in the east and *P. roemeriana* in the west, while *P. cuspidata* and *P. roemeriana* do not currently share overlapping distributions (Roda et al., 2017). These species are butterfly- and moth-pollinated (Burgin et al., 2023), and can be found growing in prairie remnants, pastures, and roadside ditches. Little is known about the specific abiotic factors distinguishing the habitats of these species; however, *P. roemeriana* has been described as specialized to more xeric and calcareous soils relative to both *P. cuspidata* and *P. drummondii*, which are both found in similarly loamy habitats (Erbe and Turner, 1962; Levin 1978a). Furthermore, unlike *P. roemeriana* and *P. drummondii*, *P. cuspidata* is self-compatible and produces abundant selfed-seed in the field (Levin 1978a; Levin 1978b; Roda and Hopkins, 2018; Shahid et al., 2024). The drastic differences in evolutionary histories of these species are also reflected in their genomic data, with *P. drummondii* having the highest heterozygosity, effective population size, and overall genetic variation (Levin, 1978b; Wu et al., in prep). Despite genomic evidence of gene flow across species (Roda et al., 2017; Garner et al., 2024; Wu et al., in prep) and interspecific compatibility in greenhouse settings (Suni and Hopkins, 2018), F1s among species pairs are only documented in low frequency (Hopkins and Rausher, 2012; Hopkins et al., 2014; Wu et al., in prep). Given the overlapping ranges, close relatedness, differing evolutionary histories, and ongoing geneflow of these species, Texas annual *Phlox* provide an excellent system for analyzing the causes and consequences of trait divergence and plasticity to environmental stimuli.

**Figure 1.**
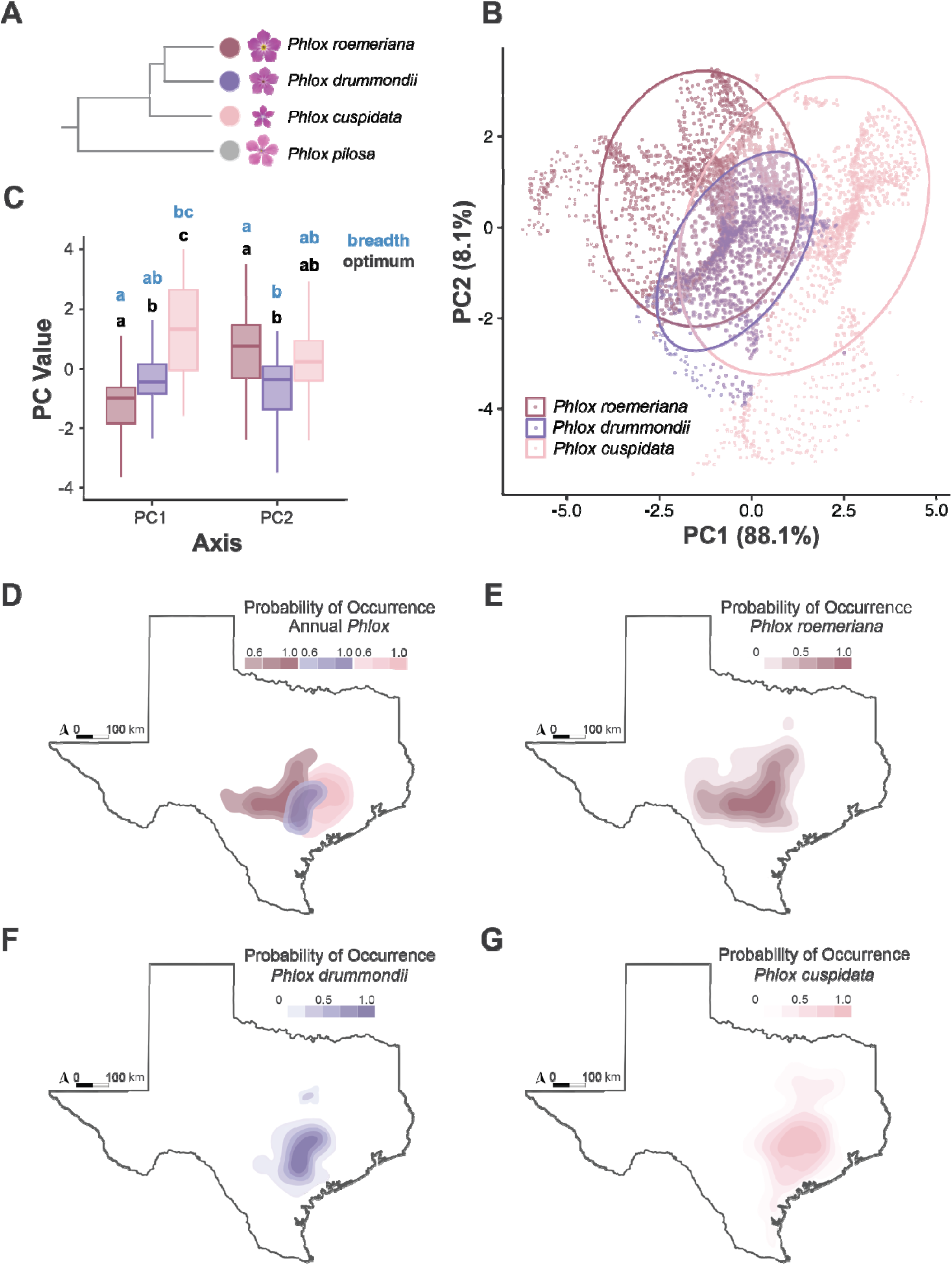
Characterization of niche for *Phlox cuspidata* (light pink), *Phlox drummondii* (purple), and *Phlox roemeriana* (dark pink). A) Schematic of the phylogenetic relationship between Texas annual *Phlox*, adapted from Garner et al., 2024. B) First two principal components of a PCA with projected distributions and WorldClim environmental variables, where each point represents a single pixel in our distribution model. C) Optimum and breadth of principle components from PCA of projected distributions. Comparisons are made across taxa, where niche optimum and breadth correspond to the median and length of the 95% inter-percentile interval along the two PCA axes respectively. Unique lettering refers to a significant difference between species in niche optimum and/or breadth, *P* < 0.05, D) Overlapping projected distributions of Texas annual *Phlox*, where probability of occurrence > 0.5, E) Projected distribution of *Phlox roemeriana*, F) Projected distribution of *Phlox drummondii*, G) Projected distribution of *Phlox cuspidata*.

Here, we use niche modeling and empirical soil testing to examine the abiotic environments of *P. cuspidata*, *P. drummondii*, and *P. roemeriana* and find that water availability is a major factor distinguishing the ranges of these species. This motivated our controlled water-availability experiment, documenting intra- and inter-species differences in plastic response to drought. We specifically interrogate whether and how differences in evolutionary history, relatedness, and biogeography influence the direction and magnitude of phenotypic response to drought.

## METHODS

### Niche modeling

We characterized the environmental niches of each of the three annual Texas *Phlox* species using species distribution modeling. All analyses were conducted in R version 4.3.3 (R Core Team, 2024). We acquired population location data from across the ranges of the focal species based on records from our recent field collections and supplemented with distribution records from the Global Biodiversity Information Facility (GBIF; downloaded 10 Dec 2023, www.Gbif.org). We filtered GBIF occurrences to include only locations with verified images. This provided 324, 180, and 296 total locations for *P. drummondii*, *P. cuspidata*, and *P. roemeriana*, respectively (N = 800). We thinned these occurrence points to one location for every five-kilometer grid cell to reduce sampling bias using the *thin* function from the ‘sp’ package version 2.1-1 (Bivand et al., 2008). We further reduced our dataset to 180 populations per species (N = 540) for consistent modeling across species. To characterize potentially uninhabitable sites, we created 1000 pseudoabsences of each species by using the *randomPoints* function from the ’dismo’ package version 1.3-14 (Hijmans and Elith, 2013).

We extracted environmental variables from the WorldClim database (downloaded 10 Dec 2023; www.WorldClim.com/version2) using the *getData* function from the ‘raster’ package version 3.6-26 (Hijmans and van Etten, 2012). Layers were trimmed and aligned to only include the state of Texas (latitude 25.5° to 35°; longitude -104° to -94°), representing an area of intermediate size that the focal taxa likely have had the opportunity to disperse into (Sobel, 2014). We selected seven layers based on their weak correlations with other layers (|r| < 0.7) and high importance to distribution models using the function *select07* from the ‘mecofun’ package version 0.5.1 (Dormann et al., 2013; Table S1). Variables were consistent among species and included three temperature and four precipitation layers at 2.5-minute resolution.

We used the *maxent* function from the ‘Maxent’ package version 3.4.0 (Phillips et al., 2006; Elith et al., 2011) to generate individual distribution models for each species. Maxent compares the distributions of environmental variables at sites occupied by a focal species to the distributions of pseudoabsences (Phillips et al., 2006; Elith et al., 2011). We used the *kfold* function from the ‘dismo’ package to cross-validate the distribution model of each species, as well as the *evaluate* function from the ‘dismo’ package to calculate the area under the receiver operator curve (AUC). The AUC is a threshold-independent indicator of species distribution model performance that ranges from 0.5 to 1, where a value of 0.5 indicates that a model is equivalent to a random draw and a value of 1 indicates that a model perfectly predicts the suitable habitat of a species (Phillips et al., 2006; Sobel, 2014).

To assess differences in habitat among species we used three separate approaches. First, we filtered the distributions of each species to include only pixels/locations of populations that had a probability of occurrence greater than 0.5. We extracted from the environmental layers from these predicted distributions of each species and used a Principal Component Analysis (PCA) to visualize the difference in habitat, niche breadth (the width of the environmental range tolerated by a species), and niche optima (the mean environmental variable value at the location with the highest predicted suitability) of each species. We then analyzed this pixel data using a one-way Multivariate Analysis of Variance (MANOVA) to construct 95% confidence intervals, as described in Sobel and Streisfeld (2015). We performed post hoc analyses on each of the seven environmental variables using the *emmeans* function from the ‘emmeans’ package version 1.10.0 (Lenth et al., 2024) to determine pairwise differences in species distributions. Finally, we used the *nicheOverlap* function from the ‘dismo’ package on pairs of species to determine Schoener’s D, quantifying the extent to which a pair of species may interact in the same space. Schoener’s D ranges from 0 to 1, with a value of 0 indicating no overlap in predicted species distributions and a value of 1 indicating complete overlap in predicted species distributions (Schoener, 1968).

### Soil testing

We characterized properties of the soil in which the three *Phlox* species grow using samples collected over a single week in May 2023 from across the range of each taxon. To sample soil at each location, we first cleared surface litter from the site and estimated soil moisture in parts per million with a moisture probe (Atree Soil Tester, Model no. B0DRFYCTYG). We then used a spade to dig a 6-inch-deep V-shaped hole and collected a 1-inch x 1-inch core from the spade. We collected cores from 10-15 sites within each location to create a population composite sample in a 1-pint plastic bag. We collected composite samples at five locations of each species (N = 15 composite samples) and left each bag open for at least 24 hours to air dry. We sent samples to the Soil, Water, and Forage Testing Laboratory at Texas A&M AgriLife (College Station, TX) to estimate a suite of chemical and physical traits including micro and macro nutrients, proportion of organic matter, and particle composition. For each variable measured, we ran an Analysis of Variance (ANOVA) test comparing the three species, using base R’s *oneway.test* function. We also used the *cor* function to determine the Pearson correlation coefficient between each variable. Finally, we performed a PCA to examine patterns of overall differentiation among species in soil properties.

### Drydown experiment

We performed a controlled dry-down experiment to characterize growth habit of the three Texas *Phlox* species and their hybrids in well-watered and dry conditions, and the differences in plasticity – response to drying – across the clade.

#### Experimental seeds

Seeds of each species were collected across their native ranges in Texas in May 2019. We used seeds from 10 *P. drummondii*, 8 *P. cuspidata*, and 4 *P. roemeriana* populations (Table S2). The wild-collected seeds were grown under controlled conditions in the greenhouses at the Arnold Arboretum of Harvard University (Boston, MA, USA) and used to generate experimental seeds from half-sib families using controlled crosses. Crosses were performed within species to generate pure species plants and between species to generate hybrid seeds. To induce germination, all seeds were soaked in 500ppm Gibberellic Acid for 48 hours and cold stratified for 7 days.

Germination occurred in seedling trays filled with LM-GPS Germination Mix (Lambert, Quebec, Canada) seedling potting media in controlled growth chamber conditions with 16 hours of supplemental light at 23°C and nighttime temperature of 18°C.

#### Wet and dry treatments

We transplanted 361 germinated seedlings (1-16 individuals from each seed family from each species) two weeks after germination into 4-inch square pots containing 200-300g of Promix High Porosity Mycorrhizae soil. Due to low germination success our samples sizes were limited such that *P. drummondii* had 23 seed families with 133 individuals; *P. cuspidata* had 22 seed families with 92 individuals; *P. roemeriana* had 14 seed families with 71 individuals; *P. cuspidata-P. roemeriana* hybrid had 8 seed families with 24 individuals; *P. drummondii-P. cuspidata* hybrid had 6 seed families with 35 individuals, and *P. drummondii-P. roemeriana* hybrid had 1 seed family with 6 individuals. Plants were fully randomized and distributed across 38 trays in the greenhouse. Individuals were allowed to acclimate for one week with a standard watering regime before being assigned to a treatment. Within each seed family, siblings were split into the dry and wet treatment groups to ensure genetic representation across the two treatments (N = 177 in dry, 183 in wet).

Wet and dry soil treatments were enforced by controlling the percent soil saturation of each pot. Wet treatment pots were maintained at 70% soil moisture whereas dry treatment pots were maintained at 12% soil moisture (percentages modified from Suni et al., 2022 due to humidity conditions in the greenhouse). Previous trials with *Phlox* plants indicates that 70% soil saturation corresponds to well-watered pots and healthy flourishing plants, whereas 12% soil moisture results in plants that can survive but appear water stressed. Both treatments consisted of weighing and watering individual pots three times each week to reach target soil saturation levels. Soil saturation for each pot was calculated as *target weight = (wet weight – dry weight) * target saturation percentage + dry weight* where dry and wet weight of each pot was measured at the start of the experiment prior to transplanting plants into pots. Dry weight was defined as the weight in grams of a pot filled with dry potting media, and wet weight was defined as the weight of the pot with potting media that is saturated with water. Individuals and their pots were weighed and watered according to their assigned treatment for five weeks.

For each individual, we counted the days from treatment to the opening of the first flower and collected four flowers within 10 days of first flowering. We scanned (EPSON V600) each flower to measure mean flower tube length and mean flower area in ImageJ (Schneider et al., 2012; www.ImageJ.net). At nine weeks after germination, individuals were harvested. Upon harvesting, height, the longest leaf length, leaf count, and flower count were measured. We estimated above ground wet biomass in grams by harvesting all stem, leaf, floral and fruit tissue growing above the surface of the potting media and weighing them on a balance. We estimated dry biomass in grams by placing the tissue in a drying oven for three days before weighing it again.

#### Analyses

We summarized the phenotypic variation across all the traits using a PCA on the pure species in Scikit-learn. We used the PCA loadings to determine which traits explained the variation along the dominant axes.

We performed a dimensionality reduction on the nine phenotypic traits for the three focal species in Python version 3.11.6 (Van Rossum and Drake, 2009), using the package umap-learn version 0.5.5 (McInnes et al., 2018). To do this, a standard scaler was used on the data from the package scikit-learn version 1.3.0 (Pedregosa et al., 2011).

To quantify the response for each trait of the species to water limitation we performed a mixed-effect linear model with the function *lmer* from the package ‘lme4’ version 1.1.34 (Bates et al., 2015). We tested for an effect of species (excluding hybrids due to small sample sizes), wet/dry treatment, and the interaction across the nine phenotypic traits. Treatment, species, and their interaction were treated as fixed effects, while block effects and maternal family ID were included in the models as random effects. Paternal effects were excluded as a random effect, as it was found to be redundant with maternal family ID and led to overfitting. The fixed effects of each model were analyzed using an ANOVA test from the package ’car’ version 3.1.2 (Fox and Weisberg, 2019) with Bonferroni correction, followed by a post-hoc Tukey HSD test.

For all species and hybrids, we calculated normalized mean values for each trait by converting raw values to z-scores to facilitate comparison across traits and groups.

## RESULTS

### Niche modeling

Our species distribution models predict the suitable habitats of the three species, as indicated by high AUC values (>0.95; Table S3). These models reveal that the three taxa occupy geographically overlapping but distinct habitats (Figure 1B-G) across an east to west gradient, with *P. cuspidata* predicted to occur in the wettest and warmest habitats, *P. drummondii* occupying a moderate habitat, and *P. roemeriana* predicted to grow in the driest and coolest habitats (Table 1; Table S4). Overall, the species have significant differences in both niche optima and niche breadth along PC1 and PC2 (Figure 1B; Figure 1C; Table 1). In fact, each species pair inhabits significantly different conditions across each environmental variable used in our species distribution modeling (Figure S1; Table S5; Table S6). The values of pairwise Schoener’s D reflected the same trend, with the distributions of *P. drummondii* and *P. cuspidata* overlapping the most (0.77) and *P. roemeriana* and *P. cuspidata* overlapping the least (0.56). When we identified the most likely habitat of each species and isolated the WorldClim environmental variables associated with each of the species’ occupied space pixels, we found that environmental variables related to precipitation (precipitation of the driest quarter, precipitation of the coldest quarter, and precipitation of the wettest quarter) explained 88.1% of the variation in PC1. Thus, soil moisture distinguishes the species’ distributions (Figure 1C; Table S7).

**Table 1.**
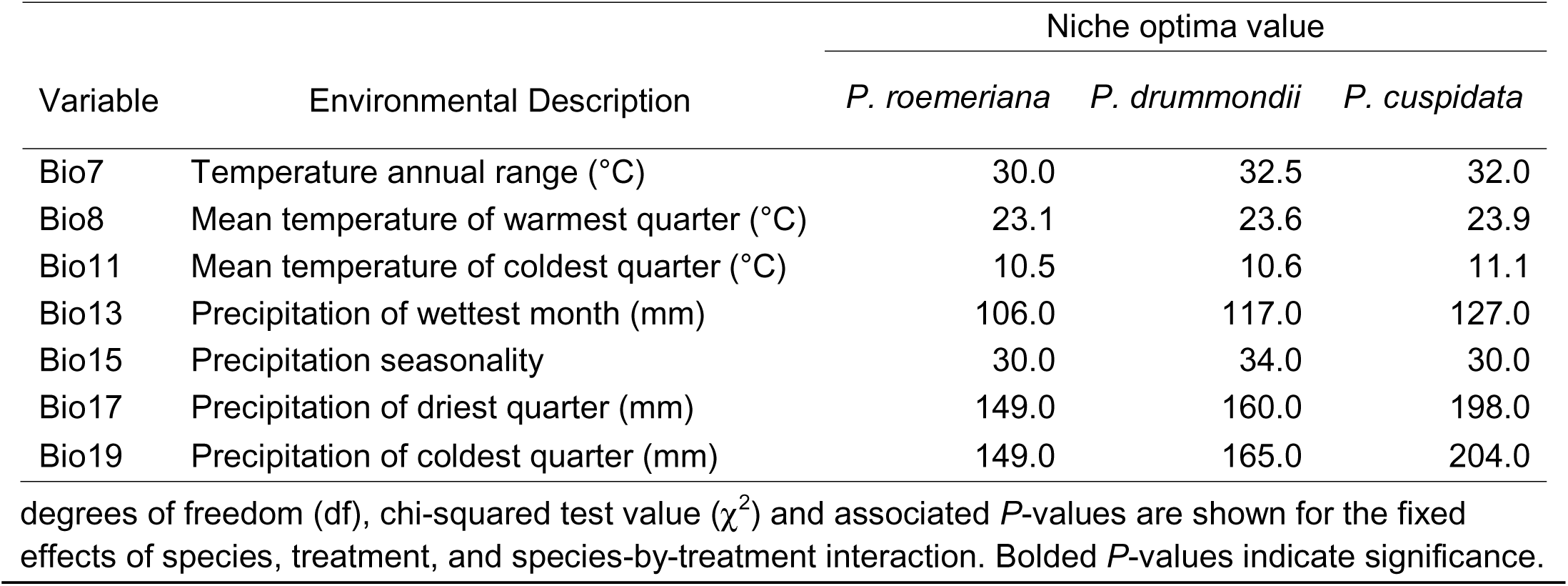
Niche optima of each species, where values represent the means of WorldClim variables in cations with the highest predicted suitability.

### Soil testing

We found substantial variation across the soil properties across species (Figure 2A-C). Each sampled metric showed significant differences across species (*p* < Bonferroni-corrected p-value < 0.05; Table S8). Soil moisture was also significantly different between the three species (Figure 2B; Table S8) with *P. cuspidata*’s environment containing higher moisture than that of *P. roemeriana* (*p <* 0.001) and *P. roemeriana*’s environment containing higher moisture than that of *P. drummondii* (*p <* 0.001). Many of the soil parameters were highly correlated with one another (Table S9), and strongly species specific, as evidenced by tight species clustering in the PCA (Figure 2C; Table S10).

**Figure 2.**
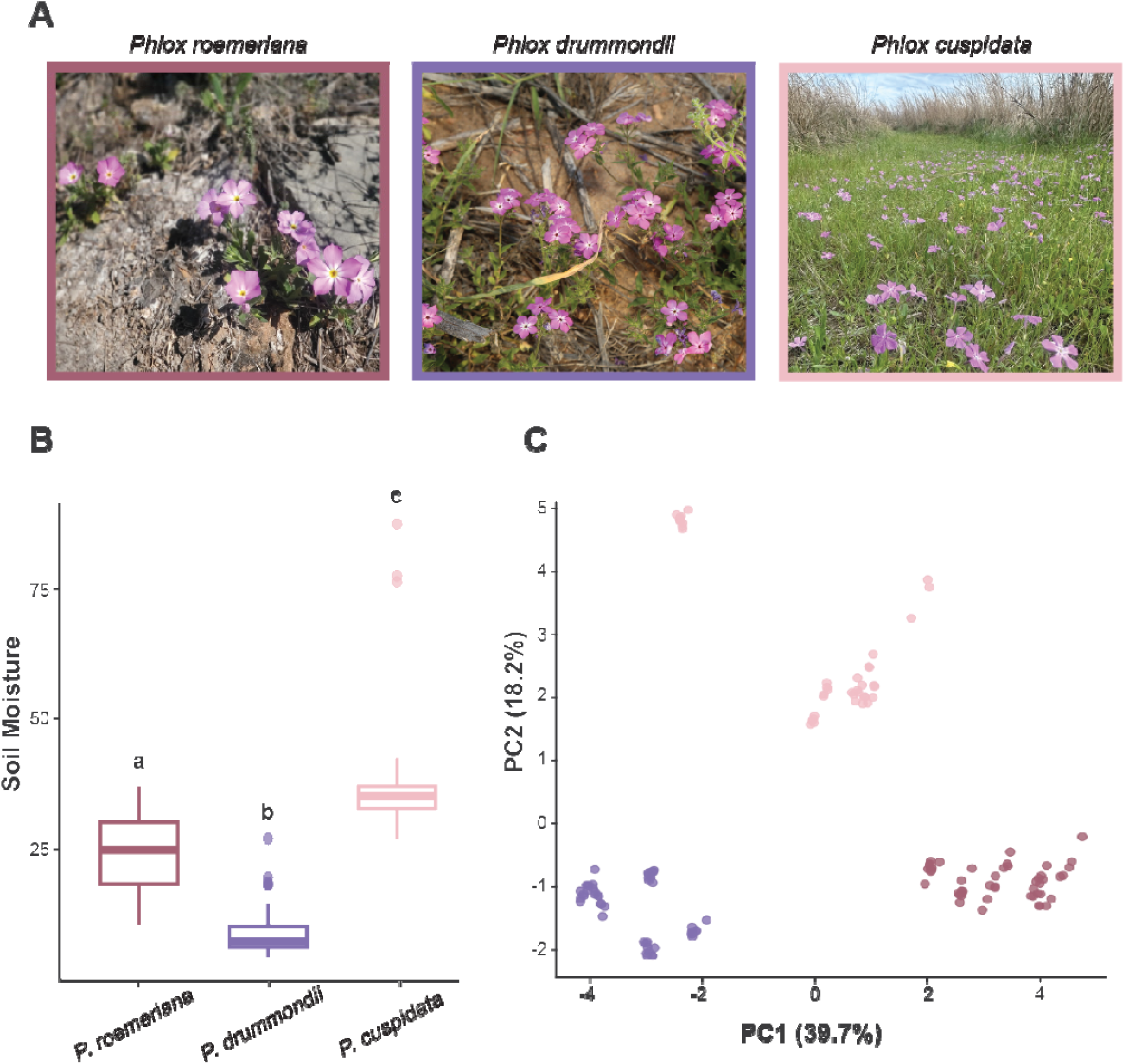
Differences in soil characteristics of *Phlox cuspidata* (light pink), *Phlox drummondii* (purple), and *Phlox roemeriana* (dark pink). A) Photographs depicting typical habitats of *P. roemeriana* (left), *P. drummondii* (center), and *P. cuspidata* (right). Photos by Patrick Alexander (https://www.inaturalist.org/observations/265914863), Victor L. Manuel (https://www.inaturalist.org/observations/281151460), and Iuisesc (https://www.inaturalist.org/observations/270333215) via iNaturalist. Licensed under CC BY-NC 4.0. B) Differences in moisture (ppm) of soil samples from natural habitats. Comparisons were done using an ANOVA test with Bonferroni correction (unique lettering indicates p < 0.05). C) Plot of the first two principal components for *P. cuspidata*, *P. drummondii*, and *P. roemeriana* based on soil samples, colored by species. See supplemental Table 10 for PCA loadings.

### Drydown experiment

To assess response to water availability and soil moisture in Texas *Phlox*, we performed a controlled drydown experiment where each species was grown in both dry and wet conditions. We evaluated constitutive differences between species as well as responses of each species to variation in conditions using both multivariate and univariate analyses of phenotypic measurements.

#### Multivariate response to water availability across species

The three Texas annual *Phlox* species have distinct phenotypes, as evidenced by both the UMAP dimensionality reduction (Figure 3A) and the results from the principal component analysis (Figure 3B). The PCA loadings indicate that the primary contributors to PC1 are mean corolla length, height, mean leaf length, fresh and dry biomass (Figure 3C; Table S11). These traits tend to reflect size and vegetative mass of a plant. *P. cuspidata* is the smallest of the three species with narrow leaves, small flowers, and short stature, while *P. drummondii* is the tallest with wide leaves, and a robust growth form. PC2 is explained by variation in flower area and days to flower and is negatively associated with flower count (Figure 3C; Table S11). These traits tend to reflect floral investment associated with sexual reproduction. *P. roemeriana* has characteristically larger flowers and a distinct inflorescence structure with fewer flowers per inflorescence and therefore it is unsurprising that this species separates from the other two species along PC2.

**Figure 3.**
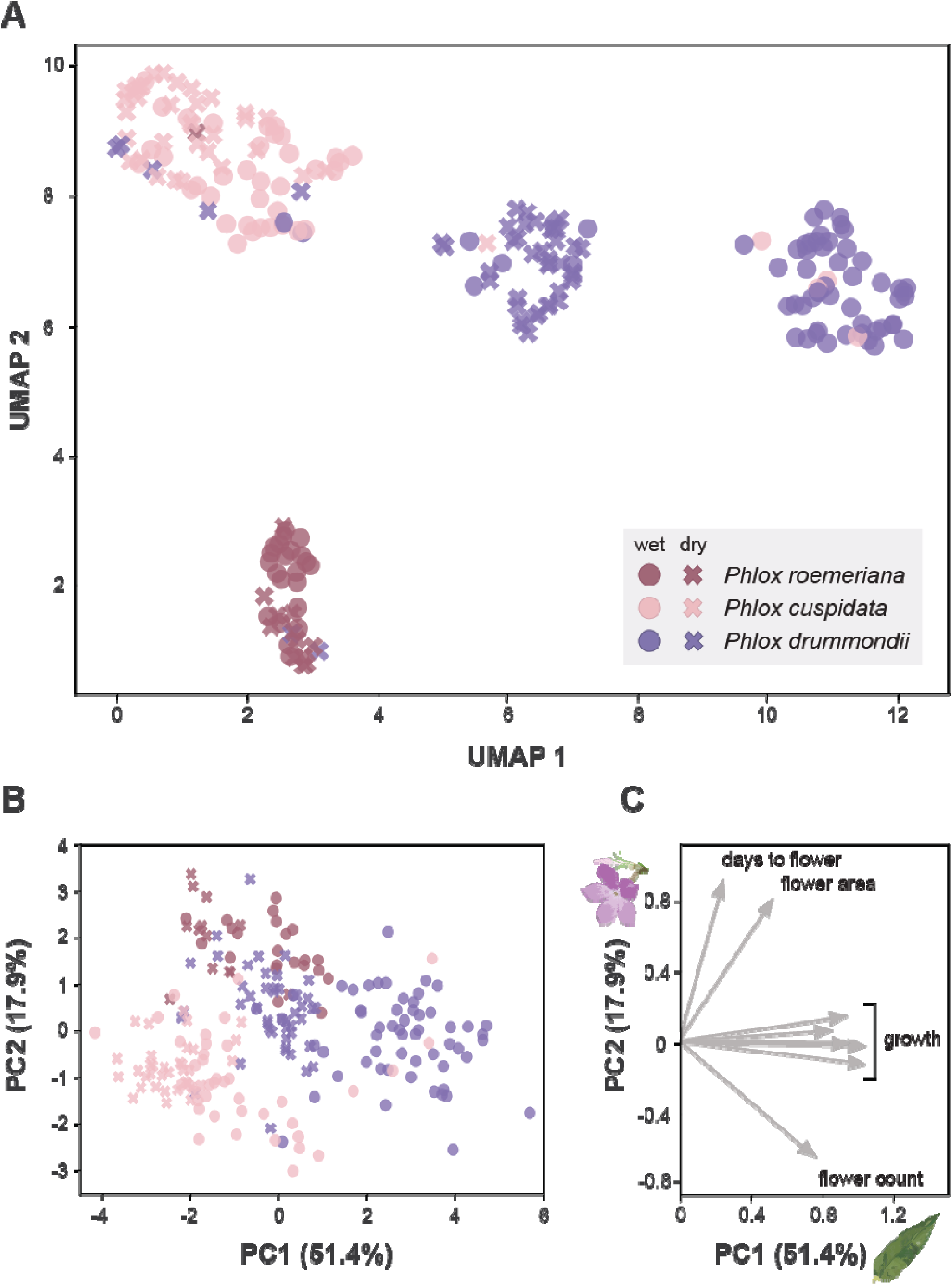
Multivariate responses to drydown experiment of annual *Phlox* species. A) UMAP plot of *Phlox cuspidata*, *Phlox drummondii*, and *Phlox roemeriana* individuals based on measurements taken during the drydown experiment, colored by species, and marked by treatment. Dimensionality reduction was performed on mean corolla length, mean flower area, height, mean leaf length, leaf count, biomass (fresh and dry), days to flower, and flower count. B) Plot of the first two principal components for *P. drummondii*, *P. cuspidata*, and *P. roemeriana* individuals based on measurements taken during the drydown experiment, colored by species, and marked by treatment. C) PCA loading arrows for all drydown variables with a loading >|0.5| for PC1 and PC2. PC1 tends to reflect overall size and vegetative investment and PC2 tends to reflect floral investment (Table S10).

Although all the species respond to water treatment, *P. drummondii* individuals respond with the greatest plasticity. This is observed in the UMAP dimensionality reduction on phenotypic traits which shows that *P. drummondii* has separate wet and dry clusters. In the PCA, the wet environment plants tend to have higher PC1 values compared to the dry environment plants for each species and this pattern is particularly robust for *P. drummondii* (Figure 3B).

#### Univariate response to water availability across species

Univariate analyses of phenotypic traits reveal broad differences between species, between treatments, and significant differences in the strength of reaction to the treatments between the three species (treatment by species interaction) (Figure 4A). Nearly all traits varied either between species, treatment, or both with only leaf count showing no significant differences. Across treatments, the rank order of species for each trait does not change (Figure 4A-B). In general, *P. drummondii* is taller, has longer leaves, more biomass, and longer corollas than the other two species. The three species differ significantly in flower area with *P. roemeriana* having the largest flowers and *P. cuspidata* having the smallest flowers.

**Figure 4.**
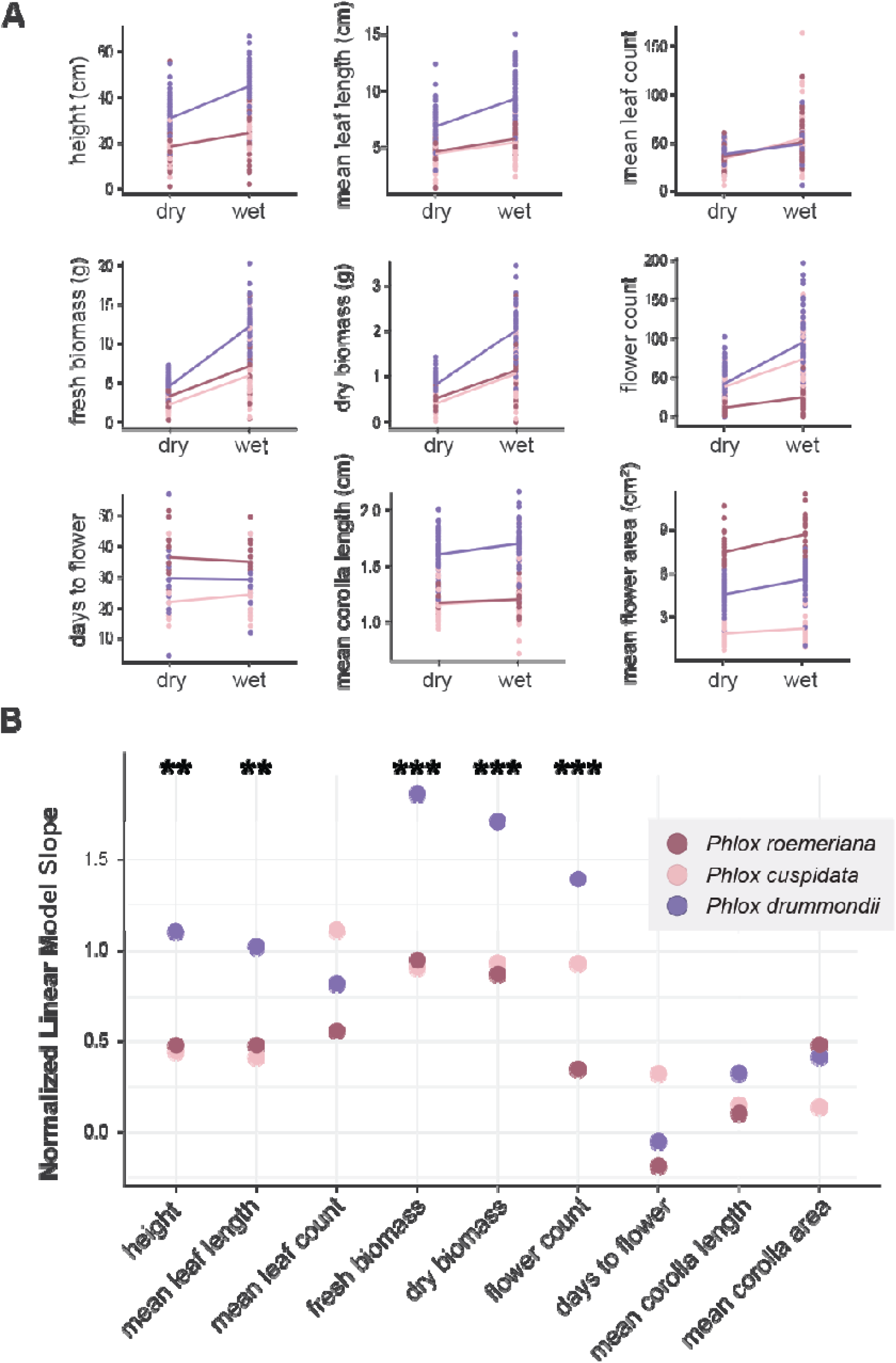
Univariate responses to drydown experiment of annual *Phlox* species. A) Reaction norms for the drydown experiment, separated by treatment and species. Lines are from a simple linear regression and only present for visualization, not analysis. B) Slopes of the linear mixed models for each species and measured variable. Data was normalized by z-score prior to plotting, in order to ensure comparable slopes. Stars indicate the significance of the species by treatment interaction, calculated with an ANOVA test on the linear mixed models with Bonferroni correction (*: p < 0.05, **: p < 0.01, ***: p < 0.001).

While the species are generally bigger with more flowers under wet conditions, the magnitude of this response is different across species as indicated by significant species by treatment interactions (p<0.05; ANOVA on linear mixed models with Bonferroni correction) for height, mean leaf length, fresh biomass, dry biomass, and flower count (Figure 4A; Figure 4B; Table 2; Table 3; Figure S2; Table S12). For all traits where the species responded differently to wet and dry conditions, *P. drummondii* has a steeper reaction norm slope than *P. cuspidata* or *P. roemeriana* (Figure 4B; Table 3; Table S12). This indicates that, while the trait measurements for *P. cuspidata* and *P. roemeriana* do vary between the wet and dry conditions, plastic response is greater in *P. drummondii.* In general, the three species tend to have more similar trait values under dry conditions with *P. drummondii* showing the greatest growth response to abundant water availability.

**Table 2.** Analysis of Variance (ANOVA) results from linear mixed models testing fixed effects of species, treatment, and their interaction on morphological and phenological traits. For each response variable, the

| Trait | Species |  |  | Treatment |  |  | Species-Treatment |  |  |
| --- | --- | --- | --- | --- | --- | --- | --- | --- | --- |
| | df | $\chi^2$ | Adj. *P* | df | $\chi^2$ | Adj. *P* | df | $\chi^2$ | Adj. *P* |
| Height | 2 | 25.014 | **< 0.001** | 1 | 79.275 | **< 0.001** | 2 | 17.772 | **0.001** |
| Leaf length | 2 | 35.305 | **< 0.001** | 1 | 63.998 | **< 0.001** | 2 | 13.756 | **0.009** |
| Leaf count | 2 | 0.204 | 1 | 1 | 61.9 | **< 0.001** | 2 | 3.9544 | 1 |
| Fresh biomass | 2 | 92.108 | **< 0.001** | 1 | 339.046 | **< 0.001** | 2 | 50.569 | **< 0.001** |
| Dry biomass | 2 | 58.199 | **< 0.001** | 1 | 245.256 | **< 0.001** | 2 | 29.914 | **< 0.001** |
| Flower count | 2 | 93.739 | **< 0.001** | 1 | 115.087 | **< 0.001** | 2 | 26.349 | **< 0.001** |
| Days to flower | 2 | 59.2564 | **< 0.001** | 1 | 0.0787 | 1 | 2 | 4.4087 | 0.992 |
| Corolla length | 2 | 113.6758 | **< 0.001** | 1 | 5.3112 | 0.1907 | 2 | 1.5072 | 1 |
| Flower area | 2 | 171.4074 | **< 0.001** | 1 | 32.6098 | **< 0.001** | 2 | 6.9286 | 0.281 |

**Table 3.** Results of Tukey’s HSD test comparing pairwise species differences under dry and wet treatments. Estimates and *P*-values are shown for each trait rison between species pairs: *Phlox cuspidata* vs *Phlox drummondii* (C-D), *Phlox cuspidata* vs *Phlox roemeriana* (C-R), and *Phlox drummondii* vs *Phlox riana* (D-R). Bolded *P*-values indicate significance (*P* < 0.05). For the full table including standard errors, degrees of freedom, and *t-*statistics, see Table S12.

| Trait | Contrast | Dry |  | Wet |  | Contrast | Within Species |  |
| --- | --- | --- | --- | --- | --- | --- | --- | --- |
|  |  | Estimate | *P*-value | Estimate | *P*-value |  | Estimate | *P*-value |
| Height | C-D | -8.536 | 0.3479 | -16.174 | 0.17 | **Dry C-Wet C** | **-5.967** | **0.0098** |
|  | C-R | 4.681 | 0.9594 | 5.120 | 0.9154 | **Dry D-Wet D** | **-13.605** | **<.0001** |
|  | **D-R** | **13.216** | **0.045** | **21.294** | **0.0001** | Dry R-Wet R | -5.527 | 0.1224 |
| Leaf length | C-D | -2.147 | 0.0315 | -3.588 | 0.1 | Dry C-Wet C | -0.975 | 0.119 |
|  | C-R | -0.066 | 1 | -0.228 | 1 | **Dry D-Wet D** | **-2.417** | **<.0001** |
|  | D-R | 2.081 | 0.848 | 3.360 | 0.5 | Dry R-Wet R | -1.138 | 0.1258 |
| Leaf count | C-D | -5.226 | 0.9991 | -0.596 | 1 | **Dry C-Wet C** | **-21.788** | **<.0001** |
|  | C-R | -1.190 | 1 | 9.120 | 0.9529 | **Dry D-Wet D** | **-17.158** | **<.0001** |
|  | D-R | 4.036 | 1 | 9.716 | 0.9435 | Dry R-Wet R | -11.479 | 0.2449 |
| Fresh biomass | **C-D** | **-2.275** | **0.0409** | **-6.115** | **< 0.0001** | **Dry C-Wet C** | **-3.607** | **<.0001** |
|  | C-R | -0.911 | 0.9737 | -1.079 | 0.9019 | **Dry D-Wet D** | **-7.446** | **<.0001** |
|  | D-R | 1.364 | 0.6913 | **5.036** | **< 0.0001** | **Dry R-Wet R** | **-3.775** | **<.0001** |
| Dry biomass | C-D | -0.423 | 0.0532 | **-0.963** | **< 0.0001** | **Dry C-Wet C** | **-0.638** | **<.0001** |
|  | C-R | -0.094 | 0.9999 | -0.055 | 1 | **Dry D-Wet D** | **-1.178** | **<.0001** |
|  | D-R | 0.329 | 0.3813 | **0.908** | **< 0.0001** | **Dry R-Wet R** | **-0.599** | **<.0001** |
| Flower count | C-D | -5.381 | 0.9989 | **-22.959** | **0.0315** | **Dry C-Wet C** | **-35.019** | **<.0001** |
|  | **C-R** | **25.541** | **0.0304** | **47.414** | **< 0.0001** | **Dry D-Wet D** | **-52.598** | **<.0001** |
|  | **D-R** | **30.922** | **0.0031** | **70.373** | **< 0.0001** | Dry R-Wet R | -13.146 | 0.62 |
| Days to flower | **C-D** | **-6.405** | **0.0093** | -3.605 | 0.443 | Dry C-Wet C | -2.392 | 0.6891 |
|  | **C-R** | **-12.846** | **< 0.0001** | **-9.065** | **0.0002** | Dry D-Wet D | 0.409 | 1 |
|  | **D-R** | **-6.440** | **0.0222** | -5.461 | 0.0836 | Dry R-Wet R | 1.389 | 0.9985 |
| Corolla length | **C-D** | **-0.413** | **< 0.0001** | **-0.460** | **< 0.0001** | Dry C-Wet C | -0.044 | 0.9843 |
|  | C-R | -0.008 | 1 | 0.004 | 1 | Dry D-Wet D | -0.091 | 0.1232 |
|  | **D-R** | **0.405** | **< 0.0001** | **0.464** | **< 0.0001** | Dry R-Wet R | -0.032 | 1 |
| Flower area | **C-D** | **-1.824** | **0.0141** | **-2.643** | **0.0001** | Dry C-Wet C | -0.309 | 0.9911 |
|  | **C-R** | **-4.882** | **< 0.0001** | **-5.786** | **< 0.0001** | **Dry D-Wet D** | **-1.128** | **0.0001** |
|  | **D-R** | **-3.058** | **0.0001** | **-3.143** | **< 0.0001** | **Dry R-Wet R** | **-1.213** | **0.0141** |

Given the differential responses of the three species to water limitation, we next looked at the responses of hybrids between the species. Interestingly, the two species that tended to be the most similar and have the weakest response to the treatment, *P. cuspidata* and *P. roemeriana*, had transgressive phenotypes in their hybrids. *P. cuspidata*-*P. roemeriana* hybrids were larger than their parents in most traits (Figure 5A; Table 2; Table 3; Table S12), particularly for the aforementioned vegetative growth traits (Hoover et al., 2017; Gupta et al., 2020). Trait values are particularly transgressive in the wet condition. In contrast, *P. drummondii*-*P. roemeriana* and *P. cuspidata*-*P. drummondii* hybrids are consistently intermediate between that of their parents (Figure 5B & 5C), a trend that persisted in both wet and dry conditions. When a principal component analysis was performed and the traits were not taken in isolation, the F1 hybrids of each of the species similarly fell in between their parents along the first two PCs (Figure S3), especially in the wet condition. In sum, the hybrids with *P. drummondii* parentage have consistent intermediate phenotype in contrast to the transgressive phenotypes seen in *P. cuspidata-P. roemeriana* hybrids.

**Figure 5.**
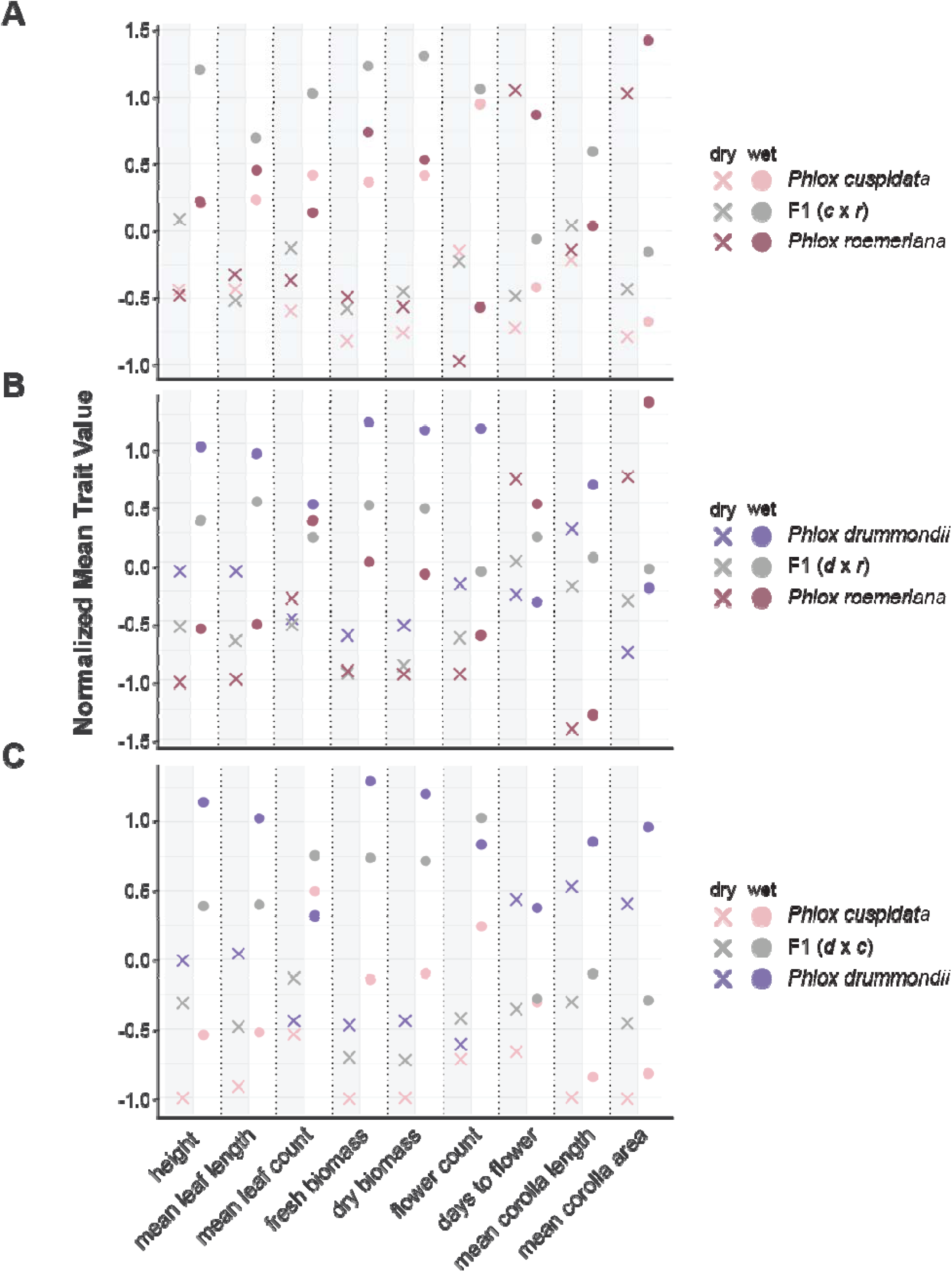
Differences in trait means of parental species and pairwise hybrids in response to drought. A) Mean trait values for *P. drummondii*, *P. roemeriana*, and their F1 hybrids in wet and dry conditions. B) Mean trait values for *P. cuspidata*, *P. roemeriana*, and their F1 hybrids. C) Mean trait values for *P. cuspidata*, *P. drummondii*, and their F1 hybrids. All data z-score adjusted.

## Discussion

We find that this three species clade of annual *Phlox* species native to the hot and dry environments of Texas occupies distinct ecological niches and expresses divergent phenotypes, particularly in response to water availability. Despite their close evolutionary relationship, these species differ in both their environmental associations and their patterns of phenotypic plasticity.

The different suitable niches inferred for each of the species align with described ecoregions in Texas along a similar east-west gradient from the Gulf Coast Prairies and Marshes to the East Central Texas Prairies, to the Edwards Plateau (Griffith et al., 2004). Ecological differentiation among the three species is primarily driven by variation in precipitation patterns and soil moisture. The major axis of environmental variation (PC1; Figure 1B; Figure 1C; Table 1) distinguishes species based on water availability, with *P. roemeriana* occupying the coolest, driest habitats, *P. cuspidata* restricted to warmer, wetter sites, and *P. drummondii* occurring in intermediate habitats. Notably, soil properties likely interact with precipitation to influence soil moisture availability, with *P. drummondii* habitats exhibiting the lowest soil moisture despite intermediate precipitation levels (Figure 2A-C; Table S10). The breadth of environmental niche also varies, with the highly selfing *P. cuspidata* maintaining the broadest niche and the limestone specialist *P. roemeriana* occupying the narrowest niche along PC1 (Figure 1B; Figure 1C; Table 1).

In the case of these three *Phlox*, species trait differences do not align neatly with expectations based on soil water availability. It has been noted that plants adapted to low water availability can have small leaf area, less biomass, and earlier flowering as part of a drought escape strategy (Chaves et al., 2003; Exposito-Alonso et al., 2018) The species with the lowest biomass, narrowest leaves, and smallest flowers, earliest flowering time is *P. cuspidata* (Figure 4A), which also occurs in the wettest soil with the highest precipitation (Table 1; Figure 2B). Contrastingly, *P. drummondii* has the most substantial biomass, biggest leaves, and tall stature, despite occupying drier habitats with the driest soils (Table 3; Table S12). Though *P. roemeriana* occurs in the hottest sites with the lowest levels of precipitation (Table 1), it produces large flowers with a distinctive inflorescence structure but intermediate vegetative size (Figure 4A). These findings suggest that vegetative traits may be shaped by selective pressures other than water availability alone, including nutrient availability linked to soil properties, or differences in life-history strategies such as selfing versus outcrossing. In this way, early flowering, and reduced size of *P. cuspidata* may reflect selection for rapid life cycle completion in ephemeral wet environments or genetic constraints imposed by its selfing mating system.

All three species exhibited substantial phenotypic plasticity in response to environmental variation in water availability. Consistently, all three species have reduced vegetative size, biomass, and leaf production under drought (Figure 4A). However, floral traits showed limited plasticity: neither flowering time nor corolla tube length change significantly across treatments. This decoupling of vegetative and floral trait plasticity mirrors patterns reported in other systems (e.g., see Moyers et al., 2021), where reproductive traits often show evolutionary or developmental canalization relative to vegetative growth in stressful environments.

This response of *Phlox* to limited water availability is consistent with expectations for plants to reduce biomass and leaf size due to stress. Across the three species we find plasticity corresponds to a major shift in morphological space from wet to dry along the growth axis PC1 and UMAP1 (Figure 3A-B). This axis also corresponds to significant differentiation between the three species but does not correspond to the associated niche differentiation inferred between the species. Specifically, *P. cuspidata* occupies the wettest habitat and yet has the most negative values along PC1 in morphological space which is associated with a plastic response to dry conditions, while *P. drummondii* occupies the driest soils in its native range but morphologically occupies the most positive PC1 values corresponding to well-watered phenotypes (Figure 3A-B). This pattern suggests that the direction of phenotypic plasticity does not match the direction of phenotypic divergence. Instead, this pattern could further indicate that broad patterns of vegetative and morphological divergence between the species are not predominantly driven by adaptation to water availability. In the case of *Phlox,* this plastic response did not correspond to a path of least resistance in ecological divergence (Fox and et al., 2019, Lei et al., 2022). Similarly, patterns of morphological divergence do not appear to be the result of canalization of a plastic response after persistent selection (as in (Agrawal et al., 1999; Chun et al., 2007; Wund et al., 2008)

Among the three species, *P. drummondii* displayed the strongest plastic response to water availability (Figure 3A-B; Figure 4A-B). Under dry conditions, species converged on similar reduced vegetative display, while *P. drummondii* displayed a disproportionate release from drought and subsequent shift in phenotype when under wet conditions relative to either *P. cuspidata* or *P. roemeriana* (Figure 3A; Figure 4A; Figure 4B). Previous work hypothesizes that plasticity may be positively correlated with environmental niche breadth (Pfenning et al., 2010; Leung et al., 2020; Stotz et al., 2021) In the case of these *Phlox*, we find that *P. drummondii* occupies an intermediate habitat, but does not inhabit the broadest environmental niche. Nevertheless, its ability to capitalize on high moisture environments suggests a flexible growth strategy constrained primarily by external water availability.

Finally, we found no evidence of hybrid dysfunction among the three species, as all F1 hybrids exhibited vegetative and floral trait values intermediate to or exceeding those of their parental species (Figure 5A-C). Notably, hybrids between *P. cuspidata* and *P. roemeriana* displayed transgressive traits, becoming larger than either parent, while hybrids with *P. drummondii* as a parent were typically intermediate (Figure 5A-C). These results show that ecological divergence among these species has not generated intrinsic barriers severe enough to limit hybrid performance, at least in terms of growth, viability, and floral morphology. This observation suggests that hybridization in this clade may not incur immediate ecological fitness costs and could contribute to ecological or phenotypic novelty in natural populations.

Our findings underscore the complex interplay between ecology, life history, and phenotypic plasticity in shaping patterns of divergence in annual *Phlox*. Future work integrating nutrient gradients, reproductive output, and fitness under natural field conditions will be crucial to fully understand the selective forces maintaining species differences and plasticity in this system.

## Acknowledgements

The authors thank Benjamin Goulet-Scott, Grace Burgin, Patrick McKenzie, and Felix Wu for guidance in developing code and interpreting results; Megan Ardolino, Scott Pedemonte, Lee Toomey, and Mike Barrett for plant husbandry during this project. Pieces of this work were presented by J.C. in his senior honors thesis, and we thank the Harvard University Hoopes Prize committee for granting it an award for outstanding scholarly research.

## Data availability statement

Data and code will be available from: Dryad doi:10.5061/dryad.x3ffbg7z4

## Literature Cited

1. Agrawal, A.A. 1999. Transgenerational induction of defenses in animals and plants. Nature 401(6748): 60–63.

2. Agrawal, A.A. 2001. Phenotypic plasticity in the interactions and evolution of species. Science 294(5541): 321–326.

3. Anderson, J.T., J.H. Willis, and T. Mitchell-Olds. 2011. Evolutionary genetics of plant adaptation. Trends in Genetics 27(7): 258–266.

4. Asadyar, L., F. Fenselau de Felippes, J. Bally, C.J. Blackman, J. An, F.C. Sussmilch, L. Moghaddam, B. Williams, S.J. Blanksby, T.J. Brodribb, and P.M. Waterhouse. 2024. Evidence for within-species transition between drought response strategies in *Nicotiana benthamiana*. New Phytologist 244(2): 464–476.

5. Bartlett, M.K., T. Klein, S. Jansen, B. Choat, and L. Sack. 2016. The correlations and sequence of plant stomatal, hydraulic, and wilting responses to drought. PNAS: 113: 13098–13103.

6. Bates, D., M. Mächler, B. Bolker, and S. Walker. 2015. Fitting linear mixed-effects models uning lme4. Journal of Statistical Software 64(1): 1–48.

7. Bivand, R.S., E.J. Pebesma, V. Gómez-Rubio, and E.J. Pebesma. 2008. Applied spatial data analysis with R. Springer.

8. Bohnert, H.J., D.E. Nelson, R.G. Jensen. 1995. Adaptations to environmental stresses. The Plant Cell 7: 1099–1111.

9. Bradshaw, A.D. 1965. Evolutionary significance of phenotypic plasticity in plants. Advances in Genetics 13: 115–155.

10. Brakefield, P.M., J. Gates, D. Keys, F. Kesbeke, P.J. Wijngaarden, A. Montelro, V. French, and S.B. Carroll. 1996. Development, plasticity, and evolution of butterfly eyespot patterns. Nature 384: 236–242.

11. Burgin, G.A., O. Bronzo-Munich, A.G. Garner, I. A. Acevedo, and R. Hopkins. 2023. Characterizing each step of pollination in *Phlox drummondii* reveals that a single butterfly species predominates in the pollinator assemblage. American Journal of Botany 110(5): e16172.

12. Chapman, M.A., S.J. Hiscock, and D.A. Filatov. 2013. Genomic divergence during speciation driven by adaptation to altitude. Molecular Biology and Evolution 30: 2553–2567.

13. Chaves, M.M., J.P. Maroco, and J.S. Pereira. 2003. Understanding plant responses to drought – from genes to the whole plant. Functional Plant Biology 30: 239–264.

14. Chun, Y.J., M.L. Collyer, K.A. Moloney, and J.D. Nason. 2007. Phenotypic plasticity of native vs. invasive purple loosestrife: A two-state multivariate approach. Ecology 88(6): 1499–1512.

15. de Jong, G. 2005. Evolution of phenotypic plasticity: Patterns of plasticity and the emergence of ecotypes. New Phytologist 166: 101–117.

16. Des Marais, D.L., K.H. Hernandez, and T.E. Juenger. 2013. Genotype-by-environment interaction and plasticity: Exploring genomic responses of plants to the abiotic environment. Annual Review of Ecology, Evolution, and Systematics 44: 5–29.

17. De Witt, T.J., and S.M. Scheiner. 2004. Phenotypic plasticity: Functional and conceptual approaches. New York, NY. Oxford University Press.

18. Dittrich-Reed, D.R., and B.M. Fitzpatrick. 2012. Transgressive hybrids as hopeful monsters. Evolutionary Biology 40(2): 310–315.

19. Dobzhansky, T. 1937. Genetics and the Origin of Species. New York, NY. Columbia University Press.

20. Dormann, C.F., J. Elith, S. Bacher, C. Buchmann, G. Carl, G. Carré, J.R. García Marquéz, B. Gruber, B. Lafourcade, P.J. Leitão, T. Münkemüller, C. McClean, P.E. Obsorne, B. Reineking, B. Schröder, A.K. Skidmore, D. Zurell, and S. Lautenbach. 2013. Collinearity: a review of methods to deal with it and a simulation study evaluating their performance. Ecography 36(1): 27–46.

21. Eckstein, R.L. 2005. Differential effects of interspecific interactions and water availability on survival, growth, and fecundity of three congeneric grassland herbs. New Phytologist 166(2): 525–536.

22. El Hafid, R., D.H. Smith, M. Karrou, and K. Samir. 1998. Physiological responses of spring durum whwat cultivars to early-season drought in a Mediterranean environment. Annals of Botany 81: 363–370.

23. Elith, J., S.J. Phillips, T. Hastie, M. Dudík, Y.E. Chee, and C.J. Yates. 2011. A statistical explanation of Maxent for ecologists. Diversity and Distributions 17: 43–57.

24. Exposito-Alonso, M., F. Vasseur, W. Ding, G. Wang, H.A. Burbano, and D. Weigel. 2018. Genomic basis and evolutionary potential for extreme drought adaptation in *Arabidopsis thaliana*. Nature Ecology & Evolution 2: 352–358.

25. Fox J., and Weisberg S. 2019. An R Companion to Applied Regression, Third edition. Sage, Thousand Oaks CA.

26. Fox, R.J., J.M. Donelson, C. Schunter, T. Ravasi, and J.D. Gaitán-Espitia. 2019. Beyond buying time: The role of plasticity in phenotypic adaptation to rapid environmental change. Philosophical Transactions of the Royal Society B: Biological Sciences 374(1768): e20180714.

27. Freedman, M.G., H. Dingle, C.A. Tabuloc, J. C. Chiu, L.H. Pang, and M.P. Zalucki. 2017. Non-migratory monarch butterflies, *Danaus plexippus* (L.), retain developmental plasticity and a navigational mechanism associated with migration. Biological Journal of the Linnean Society 123(2): 2018.

28. Garner, A.G., B.E. Goulet-Scott, and R. Hopkins. 2024. Phylogenomic analyses re-examine the evolution of reinforcement and hypothesized hybrid speciation in *Phlox* wildflowers. New Phytologist 243(1): 451–465.

29. Ghalambor, C.K., J.K. McKay, S.P. Carroll, and D.N. Reznick. 2007. Adaptive versus non-adaptive phenotypic plasticity and the potential for contemporary adaptation in new environments. Functional Ecology 21(3): 394–407.

30. Gibert, J. 2017. The flexible stem hypothesis: Evidence from genetic data. Developmental Genes and Ecology 227: 297–307.

31. Griffith, G.E., S.A. Bryce, J.M. Omernik, J. Comstock, A. Rogers, B. Harrison, S. Hatch, and D. Bexanson. 2004. Ecoregions of Texas. USGS, Corvalis, OR.

32. Gupta, A., A. Rico-Medina, and A.I. Caño-Delgado. 2020. The physiology of plant responses to drought. Science 368(6488): 266–269.

33. Hijmans, R.J., and J. van Etten. 2012. raster: Geographic analysis and modeling with raster data. R package version 2.0–12.

34. Hijmans, R.J., and J. van Etten. 2013. Species distribution modeling with R. R CRAN Project.

35. Hoffman, A.A., and J.R. Bridle. 2021. The dangers of irreversibility in an age of increased uncertainty: Revisiting plasticity in invertebrates. Oikos 2022(4): e08715.

36. Hoover, D.L., M.C. Duniway, and J. Belnap. 2017. Testing the apparent resistance of three dominant plants to chronic drought on the Colorado Plateau. Journal of Ecology 105: 152–162.

37. Hopkins, R., and M.D. Rausher. 2012. Pollinator-mediated selection on flower color allele drives reinforcement. Science 335(6072): 1090–1092.

38. Hopkins, R., R.F. Guerrero, M.D. Rausher, and M. Kirkpatrick. 2014. Strong reinforcing selection in a Texas wildflower. Current Biology 24(17): 1995–1999.

39. Husemann, M., M. Tobler, C. McCauley, B. Ding, and P.D. Danley. 2017. Body shape differences in a pair of closely related Malawi cichlids and their hybrids: Effects of genetic variation, phenotypic plasticity, and transgressive segregation. Ecology and Evolution 7: 4336–4346.

40. Kagawa, K., and G. Takimoto. Hybridizatoin can promote adaptive radiation by means of transgressive segregation. Ecology Letters 21(2): 264–274.

41. Kawecki, T.J., and D. Ebert. 2004. Conceptual issues in local adaptation. Ecology Letters 7(12): 1225–1241.

42. Kellermann, V., A.A. Hoffmann, J. Overgaard, V. Loeschcke, and C.M. Sgrò. 2018. Plasticity for desiccation tolerance across *Drosophila* species is affected by phylogeny and climate in complex ways. Proceedings of the Royal Society B: Biological Sciences 285: e20180048.

43. Kulkarni, S.S., I. Gomez-Mestre, C.L. Moskalik, B.L. Storz, and D.R. Buchholz. 2011. Evolutionary reduction of developmental plasticity in desert spadefoot toads. Journal of Evolutionary Biology 24: 2445–2455.

44. Lenth, R.V., B. Bolker, P. Buerkner, I. Giné-Vásquez, M. Herve, M. Jung, J. Love, F. Miguez, H. Riebl, and H. Singmann. 2024. emmeans: Estimates marginal means, aka least-squares means. R package 1.10.0.

45. Levin, D.A. 1978a. Genetic variation in annual *Phlox*: Self-compatible versus self-incompatible species. Evolution 32: 245–263.

46. Levin, D.A., 1978b. Inbreeding depression in partially self-fertilizing *Phlox*. Evolution 43(7): 1417–1423.

47. Leung, C., M. Rescan, D. Grulois, and L-M. Chevin. 2020. Reduced phenotypic plasticity evolves in less predictable environments. Ecology Letters 23(11): 1664–1672.

48. Lind, M.I., K. Yarlett, J. Reger, M.J. Carter, and A.P. Beckerman. 2015. The alignment between phenotypic plasticity, the major axis of genetic variation, and the response to selection. Proceedings of the Royal Society B: Biological Sciences 282(1816): e20151651.

49. Liu, D., J. Peñuelas, R. Ogaya, M. Estiarte, K. Tielbörger, F. Slowik, and X. Yang. 2017. Species selection under long-term experimental warming and drought explained by climatic distributions. New Phytologist 217(4): 1494–1506.

50. Liu, H., Q. Ye, K.J. Simpson, E. Cui, and J. Xia. 2022. Can evolutionary history predict plant plastic responses to climate change? New Phytologist 235(3): 1260–1271.

51. Losos, J.B., T.R. Jackman, A. Larson, K. de Queiroz, and L. Rodriguez-Schettino. 1998. Contingency and determinism in replicated adaptive radiations of island lizards. Science 279(5359): 2115–2118.

52. McInnes, L., J. Healy, and J. Melville. 2018. Umap: Uniform manifold approximation and projection for dimension reduction. arXiv preprint arXiv: 1802.03426.

53. Moyers, J., and M.M. Kotowska. 2021. A meta-analysis of responses in floral traits and flower-visitations to water deficit. Global Change Biology 27(13): 3095–3108.

54. Muller, H.J. 1942. Isolating mechanisms, evolution, and temperature. Biological Symposia 6: 71–125.

55. Nielsen, M.E., and D.R. Papaj. 2022. Why study plasticity in multiple traits? New hypotheses for how phenotypically plastic traits interact during development and selection. Evolution 76(5): 858–869.

56. Pedregosa, F., G. Varoquaux, A. Gramfort, V. Michel, B. Thirion, O. Grisel, M. Blondel, P. Prettenhofer, P. Weiss, V. Dubourg, J. and Vanderplas. 2011. Scikit-learn: Machine learning in Python. The Journal of Machine Learning Research 12: 2825–2830.

57. Pfenning, D.W., M.A. Wund, E.C. Snell-Rood, T. Cruickshank, C.D. Schlichting, and A.P. Moczek. 2010. Phenotypic plasticity’s impacts on diversification and speciation. Trends in Ecology & Evolution 25: 459–467.

58. Phillips, S.J., R.P. Anderson, and R.E. Schapire. 2006. Maximum entropy modeling of species geographic distributions. Ecological Modelling 190: 231–259.

59. Pigliucci, M., K. Cammell, and J. Schmitt. 1999. Evolution of phenotypic plasticity: A comparative approach in the phylogenetic neighborhood of *Arabidopsis thaliana*. Journal of Evolutionary Biology 12: 779–791.

60. Pollard, H., M. Cruzan, and M. Pigliucci. 2001. Comparative studies of reaction norms in *Arabidopsis* I. Evolution of response to daylength. Evolutionary Ecology Research 3: 129–155.

61. R Core Team. 2024. A language and environment for statistical computing. R Foundation for Statistical Computing, Vienna, Austria.

62. Radersma, R., D.W.A. Noble, and T. Uller. 2020. Plasticity leaves a phenotypic signature during local adaptation. Evolution Letters 4: 360–370.

63. Rieseberg, L.H., M.A. Archer, and R.K. Wayne. 1999. Transgressive segregation, adaptation, and speciation. Heredity 83: 363–372.

64. Reiseberg, L.H., A. Widmer, M. Arntz, and B. Burke. 2003. The genetic architecture necessary for transgressive segregation is common in both natural and domesticated populations. Philosophical Transactions of the Royal Society B: Biological Sciences 358: 1141–1147.

65. Roff, D. 1999. Phenotypic evolution – A reaction norm perspective. Heredity 82(344): 343–345.

66. Roda, F., F.K. Mendes, M.W. Hahn, and R. Hopkins. 2017. Genomic evidence of gene flow during reinforcement in Texas *Phlox*. Molecular Ecology 26(8): 2317–2330.

67. Roda, F., and R. Hopkins. 2018. Correlated evolution of self- and interspecific incompatibility across the range of a Texas wildflower. New Phytologist 221(1): 553–564.

68. Schneider, C.A., W.S. Rasband, and K.W. Eliceiri. 2012. NIH image to ImageJ: 25 years of image analysis. Nature Methods 9(7): 671–675.

69. Schoener, T.W. 1968. The Anolis lizards of Bimini: Resource partitioning in a complex fauna. Ecology 49: 704–726.

70. Shahid, B.M., G.A. Burgin, and R. Hopkins. 2024. Experimental and genetic analyses of selfing reveals no reinforcement in *Phlox cuspidata*. International Journal of Plant Sciences 185(3): 228–237.

71. Sicard, A., A. Thamm, C. Marona, Y. Wha Lee, V. Wahl, J.R. Stinchcombe, S.I. Wright, C. Kappel, and M. Lenhard. 2014. Repeated evolutionary changes of leaf morphology caused by mutations to a homeobox gene. Current Biology 24(6): 1880–1886.

72. Sobel, J.M. 2014. Ecogeographic isolation and speciation in the genus *Mimulus*. The American Naturalist 184(5): 565–579.

73. Sobel, J.M., and M.A. Streisfeld. 2015. Strong premating reproductive isolation drives incipient speciation in *Mimulus aurantiacus*. Evolution 69: 447–461.

74. Sol, D., P. Duncan, T.M. Blackburn, P. Cassey, and L. Lefebvre. 2005. Big brains, enhanced cognition, and response of birds to novel environments. PNAS 102: 5460–5465.

75. Stelkens, R., and O. Seehausen. 2009. Genetic distance between species predicts novel trait expression in their hybrids. Evolution 63: 884–897.

76. Stotz, G.C., C. Salgado-Luarte, V.M. Escobedo, F. Valladares, and E. Gianoli. 2021. Global trends in phenotypic plasticity of plants. Ecology Letters 24(10): 2267–2281.

77. Suni, S.S., B. Ainsworth, and R. Hopkins. 2020. Local adaptation mediates floral responses to water limitation in an annual wildflower. American Journal of Botany 107(2): 209–218.

78. Turelli, M., H.A. Orr, and J.A. Coyne. 2001. Theory and speciation. Trends in Ecology and Evolution 16(7): 330–343.

79. van Kleunen, M., and M. Fischer. 2005. Constraints on the evolution of adaptive phenotypic plasticity in plants. New Phytologist 166: 49–60.

80. Van Rossum, G., and F.L. Drake. 2009. Python 3 Reference Manual. Scotts Valley, CA: CreateSpace.

81. Via, S., R. Gomulkiewicz, G. De Jong, S.M. Scheiner, C.D. Schlichting, and P.H. Van Tienderen. 1995. Adaptive phenotypic plasticity: Consensus and controversy. Trends in Ecology & Evolution 10(5): 212–217.

82. Vinton, A.C., S.L.J. Gascoigne, I. Sepil, and R. Salguero-Gómez. 2022. Plasticity’s role in adaptive evolution depends on environmental change components. Trends in Ecology and Evolution 37(12): 1067–1078.

83. Waddington, C.H. 1953. Genetic assimilation of an acquired character. Evolution 7(2): 118–126.

84. Walter, G.M., J. Clark, A. Cristaudo, D. Terranove, B. Nevado, S. Catara, M. Paunov, V. Velikova, D. Filatov, S. Cozzolino, S.J. Hiscock, and J.R. Bridle. 2022. Adaptive divergence generates distinct plastic responses in two closely related *Senecio* species. Evolution 76(6): 1229–1245.

85. West-Eberhard, M.J. 1989. Phenotypic plasticity and the origins of diversity. Annual Review of Ecology, Evolution, and Systematics 20: 249–278.

86. Whitehead, A., J.L. Roach, S. Zhang, and F. Galvez. 2011. Genomic mechanisms of evolved physiological plasticity in killifish distributed along an environmental salinity gradient. PNAS 108: 6193–6198.

87. Wu, F.L., D. Khost, P.F. McKenzie, S. Chaturvedi, G.A. Burgin, and R. Hopkins. in prep. Genome assemblies of the Texas Phlox reveal pervasive gene flow in the presence of reinforcement.

88. Wund, M.A., J.A. Baker, B. Clancy, J.L. Golub, and S.A. Foster. 2008. A test of the “flexible stem” model of evolution: Ancestral plasticity, genetic accommodation, and morphological divergence in the threespine stickleback radiation. The American Naturalist 172(4): 449–462.

