## Supplemental information for "Variation in response to water availability across *Phlox* species"

| **Table S1.** Pearson correlations coefficients among the environmental variables used in species distribution modeling across Texas, USA. Environmental variables include measures of temperature and precipitation from the WorldClim dataset. Correlations were calculated across the entire study region to assess collinearity among variables prior to model selection. | | | | | | |  |
| --- | --- | --- | --- | --- | --- | --- | --- |
|  |  | Correlation coefficients | | | | | |
| Layer | Environmental Description | Bio7 | Bio8 | Bio11 | Bio13 | Bio15 | Bio17 |
| Bio7 | Temperature annual range (°C) | 1 |  |  |  |  |  |
| Bio8 | Mean temperature of warmest quarter (°C) | -0.135 | 1 |  |  |  |  |
| Bio11 | Mean temperature of coldest quarter (°C) | -0.356 | 0.929 | 1 |  |  |  |
| Bio13 | Precipitation of wettest month (mm) | -0.719 | 0.415 | 0.550 | 1 |  |  |
| Bio15 | Precipitation seasonality | 0.135 | 0.643 | 0.512 | 0.316 | 1 |  |
| Bio17 | Precipitation of driest quarter (mm) | -8.28 | -0.034 | 0.194 | 0.630 | -0.312 | 1 |
| Bio19 | Precipitation of coldest quarter (mm) | -0.742 | -0.028 | 0.196 | 0.541 | -0.410 | 0.496 |

| **Table S2.** Population ID and location for seed families used to generate seeds for the drydown experiment. | | |
| --- | --- | --- |
|  | Population coordinates | |
| Population | Latitude | Longitude |
| ***Phlox cuspidata* populations** | | |
| 647 | 30.4806167 | -96.7606167 |
| 651 | 30.567097 | -96.213673 |
| 652 | 30.5466333 | -96.5575 |
| 666 | 29.6023035 | -96.1105307 |
| 668 | 29.680228 | -96.584439 |
| 680 | 29.7466167 | -97.4097 |
| 654 | 30.07125 | -97.09005 |
| 659 | 30.1048333 | -97.29615 |
| ***Phlox drummondii* populations** | | |
| 634 | 29.4024 | -97.7745167 |
| 640 | 29.40755 | -97.7562167 |
| 645 | 29.4161 | -97.70415 |
| 656 | 30.0852333 | -96.9161667 |
| 662 | 30.0621514 | -97.3502951 |
| 676 | 29.4689024 | -97.8737457 |
| 680 | 29.7466167 | -97.4097 |
| 690 | 30.880867 | -98.653667 |
| 691 | 30.272333 | -98.841583 |
| 692 | 30.5497 | -98.110483 |
| 654 | 30.07125 | -97.09005 |
| 659 | 30.1048333 | -97.29615 |
| ***Phlox roemeriana* populations** | | |
| 683 | 29.5822333 | -98.6926667 |
| 687 | 31.1708 | -100.5119 |
| 689 | 30.959172 | -100.457004 |
| 693 | 30.480633 | -98.1982 |

| **Table S3.** Area under the receiver operator curve (AUC) values calculated from each species’ Maxent distribution model. AUC values range from 0.5 to 1, where a value of 0.5 indicates that a model is equivalent to a random draw and a value of 1 indicates that a model perfectly predicts the suitable habitat of a species. | | | |
| --- | --- | --- | --- |
|  | *P. roemeriana* | *P. drummondii* | *P. cuspidata* |
| AUC | 0.968 | 0.994 | 0.989 |

| **Table S4.** Relative contributions of selected WorldClim environmental variables to species distribution models for *Phlox roemeriana*, *Phlox drummondii*, and *P. cuspidata*. Variable contributions (%) were calculated from each species’ Maxent model and indicate the proportion of model gain explained by each environmental factor. Variables include temperature and precipitation parameters describing annual and season climatic variation. | | | | |
| --- | --- | --- | --- | --- |
| Variable | Environmental description | Contribution to predicted niche (%) | | |
|  |  | *P. roemeriana* | *P. drummondii* | *P. cuspidata* |
| Bio7 | Temperature annual range (°C) | 11.8 | 8.5 | 0.1 |
| Bio8 | Mean temperature of warmest quarter (°C) | 12.2 | 0.3 | 1.6 |
| Bio11 | Mean temperature of coldest quarter (°C) | 22.7 | 30.2 | 25.1 |
| Bio13 | Precipitation of wettest month (mm) | 5.5 | 0.3 | 9.7 |
| Bio15 | Precipitation seasonality | 0.4 | 19.8 | 9.1 |
| Bio17 | Precipitation of driest quarter (mm) | 16.8 | 6.8 | 2.6 |
| Bio19 | Precipitation of coldest quarter (mm) | 30.6 | 34.1 | 51.9 |

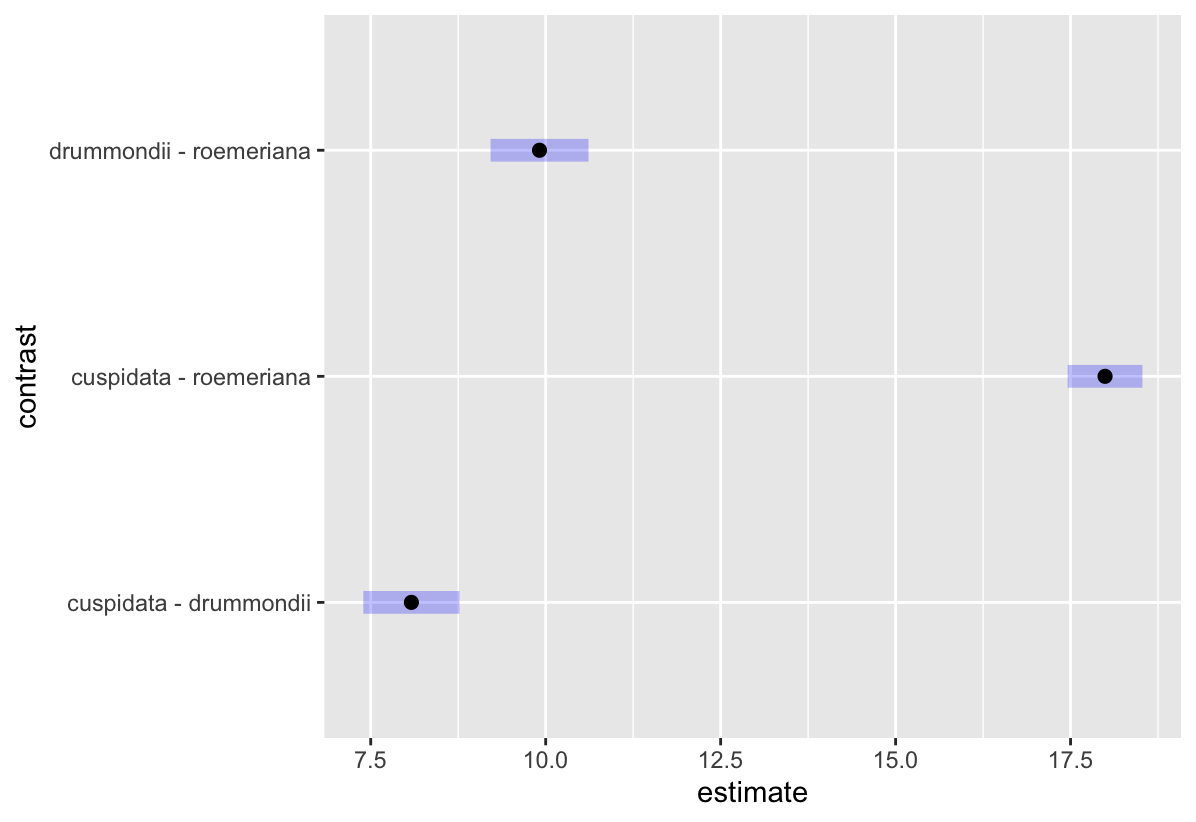

**Figure S1.** Pairwise contrasts of Maxent distribution model pixel data among *Phlox cuspidata*, *Phlox drummondii*, and *Phlox roemeriana*. Estimates represent the difference in multivariate means for each species pair, calculated from a one-way MANOVA. Error bars denote 95% confidence intervals. Larger estimates indicate greater multivariate distance in niche dimensions between species.

| **Table S5.** Summary of Multivariate Analysis of Variance (MANOVA) testing for differences in environmental conditions among *Phlox roemeriana*, *P. drummondii*, and *P. cuspidata*. Environmental variable values were extracted from predicted species distributions, and a one-way MANOVA was performed on this pixel-level data to assess multivariate differences in habitat occupancy. The significant effect of species (Pillai’s Trace = 0.847, *P* < 2.2 x 10^-16^) indicates strong multivariate differentiation in the environmental conditions associated with each species’ predicted range. | | | | | | |
| --- | --- | --- | --- | --- | --- | --- |
| Factor | df | Pillai’s Trace | Approx. *F*-value | Num. df | Den. df | *P*-value |
| Species | 2 | 0.847 | 529.94 | 14 | 10104 | **< 0.001** |
| Residuals | 5057 |  |  |  |  |  |

| **Table S6.** Results of Tukey’s HSD tests comparing pairwise species differences in environmental variable means extracted from predicted species distributions. Multiple comparisons were conducted following a one-way MANOVA on WorldClim environmental variables associated with pixel-level occurrence predictions for *Phlox cuspidata*, *Phlox drummondii*, and *Phlox cuspidata*. The table reports mean differences and adjusted *P*-values for each species pair and environmental variable. Bolded *P*-values indicate significance (*P* < 0.05) | | | |
| --- | --- | --- | --- |
| Factor | Difference | | Adjusted *P*-value |
| Bio7 |  | |  |
| *drummondii* vs. *cuspidata* | 4.51 | ***P* < 0.001** | |
| *roemeriana* vs. *cuspidata* | 9.43 | ***P* < 0.001** | |
| *roemeriana* vs. *drummondii* | 4.92 | ***P* < 0.001** | |
| Bio8 |  |  | |
| *drummondii* vs. *cuspidata* | 4.26 | ***P* < 0.001** | |
| *roemeriana* vs. *cuspidata* | -2.44 | ***P* < 0.001** | |
| *roemeriana* vs. *drummondii* | -6.70 | ***P* < 0.001** | |
| Bio11 |  |  | |
| *drummondii* vs. *cuspidata* | 1.25 | **0.0016** | |
| *roemeriana* vs. *cuspidata* | -9.16 | ***P* < 0.001** | |
| *roemeriana* vs. *drummondii* | -10.42 | ***P* < 0.001** | |
| Bio13 |  |  | |
| *drummondii* vs. *cuspidata* | -8.09 | ***P* < 0.001** | |
| *roemeriana* vs. *cuspidata* | -19.95 | ***P* < 0.001** | |
| *roemeriana* vs. *drummondii* | -11.86 | ***P* < 0.001** | |
| Bio15 |  |  | |
| *drummondii* vs. *cuspidata* | 4.50 | ***P* < 0.001** | |
| *roemeriana* vs. *cuspidata* | 5.25 | ***P* < 0.001** | |
| *roemeriana* vs. *drummondii* | 0.75 | ***P* < 0.001** | |
| Bio17 |  |  | |
| *drummondii* vs. *cuspidata* | -28.09 | ***P* < 0.001** | |
| *roemeriana* vs. *cuspidata* | -51.41 | ***P* < 0.001** | |
| *roemeriana* vs. *drummondii* | -23.31 | ***P* < 0.001** | |
| Bio19 |  |  | |
| *drummondii* vs. *cuspidata* | -33.55 | ***P* < 0.001** | |
| *roemeriana* vs. *cuspidata* | -60.83 | ***P* < 0.001** | |
| *roemeriana* vs. *drummondii* | -27.28 | ***P* < 0.001** | |

| **Table S7.** Principal Component Analysis (PCA) loadings for environmental variables associated with predicted species distributions. Loadings indicate the contribution of each WorldClim environmental variable to the first three principal components, which together explain 97.8% of the total variation in environmental niche space. Variables with higher absolute loadings contribute more strongly to each axis. | | | |
| --- | --- | --- | --- |
| Loading | PCA axis | | |
|  | PC1 (88.1%) | PC2 (8.1%) | PC3 (1.6%) |
| Bio17 | 0.636 | 0.099 | 0.150 |
| Bio11 | 0.063 | -0.623 | -0.151 |
| Bio19 | 0.730 | 0.044 | -0.111 |
| Bio7 | -0.072 | 0.582 | 0.527 |
| Bio15 | -0.074 | -0.054 | 0.081 |
| Bio13 | 0.219 | -0.117 | 0.236 |
| Bio8 | -0.017 | -0.490 | 0.776 |

| **Table S8.** Analysis of Variance (ANOVA) results for soil characteristics across sampling locations, across species. Degrees of freedom (df), sum of squares (Sum Sq), mean squares (Mean Sq), F-values, and associated *P*-values are shown for each soil parameter tested. Bolded *P*-values indicate significance (*P* < 0.05). | | | | | |
| --- | --- | --- | --- | --- | --- |
| Soil Characteristic | df | Sum Sq | Mean Sq | *F*-value | *P* -value |
| Soil moisture | 2 | 18831 | 9415 | 133.8 | **< 0.001** |
| Soil temperature | 2 | 417 | 208.66 | 7.732 | **< 0.001** |
| pH | 2 | 20.01 | 10.006 | 60.44 | **< 0.001** |
| Conductance | 2 | 9169 | 4585 | 24.06 | **< 0.001** |
| Nitrate | 2 | 1232 | 616.1 | 17.23 | **< 0.001** |
| Phosphorus | 2 | 4047 | 2023.6 | 30.57 | **< 0.001** |
| Potassium | 2 | 726263 | 363132 | 243 | **< 0.001** |
| Calcium | 2 | 5.20E+09 | 2.60E+09 | 643 | **< 0.001** |
| Magnesium | 2 | 724134 | 362067 | 148.8 | **< 0.001** |
| Sulfur | 2 | 174476 | 87238 | 566.8 | **< 0.001** |
| Sodium | 2 | 5668 | 2834 | 209.2 | **< 0.001** |
| Iron | 2 | 2629 | 1314.5 | 37.25 | **< 0.001** |
| Zinc | 2 | 18.88 | 9.439 | 9.149 | **< 0.001** |
| Manganese | 2 | 300.8 | 150.4 | 10.09 | **< 0.001** |
| Copper | 2 | 0.207 | 0.10351 | 42.41 | **< 0.001** |
| Boron | 2 | 0.6489 | 0.3245 | 40.71 | **< 0.001** |
| Percent sand | 2 | 15243 | 7622 | 515.4 | **< 0.001** |
| Percent silt | 2 | 3266 | 1633.1 | 357.4 | **< 0.001** |
| Percent clay | 2 | 5014 | 2507.1 | 296.1 | **< 0.001** |
| Percent organic carbon | 2 | 62.58 | 31.291 | 154.3 | **< 0.001** |

| **Table S9.** Pearson correlation coefficients among measured soil characteristics across sampling sites, across species. The table displays pairwise correlation values between key soil parameters, including (from top to bottom) moisture, temperature, pH, electrical conductance (EC), nutrient concentrations (nitrate, phosphorus, potassium, calcium, magnesium, sulfur, sodium), trace metals (iron, zinc, manganese, copper, boron), and soil texture variables (% sand, silt, clay, and organic carbon). These correlations reveal relationships and potential covariation patterns among soil properties relevant to plant growth and ecosystem processes. | | | | | | | | | | | |
| --- | --- | --- | --- | --- | --- | --- | --- | --- | --- | --- | --- |
|  | Correlation coefficients | | | | | | | | | | |
| Soil characteristic | Moisture | Temp | pH | EC | NON_3_ | P | K | Ca | Mg | S | Na |
| Moisture | 1 |  |  |  |  |  |  |  |  |  |  |
| Temp | 0.13 | 1 |  |  |  |  |  |  |  |  |  |
| pH | 0.10 | -0.31 | 1 |  |  |  |  |  |  |  |  |
| EC | 0.39 | -0.04 | 0.33 | 1 |  |  |  |  |  |  |  |
| NON_3_ | 0.25 | 0.11 | 0.00 | 0.85 | 1 |  |  |  |  |  |  |
| P | -0.38 | 0.18 | -0.24 | 0.03 | 0.07 | 1 |  |  |  |  |  |
| K | -0.03 | -0.28 | 0.70 | 0.05 | -0.17 | -0.17 | 1 |  |  |  |  |
| Ca | 0.22 | -0.25 | 0.73 | 0.15 | -0.09 | -0.43 | 0.83 | 1 |  |  |  |
| Mg | 0.37 | -0.08 | 0.48 | 0.10 | -0.08 | -0.46 | 0.64 | 0.66 | 1 |  |  |
| S | 0.24 | -0.24 | 0.74 | 0.18 | -0.07 | -0.43 | 0.83 | 1.00 | 0.67 | 1 |  |
| Na | 0.69 | 0.31 | -0.38 | 0.06 | 0.04 | -0.38 | -0.36 | -0.12 | 0.30 | -0.10 | 1 |
| Fe | 0.18 | 0.16 | -0.82 | -0.10 | 0.05 | -0.12 | -0.56 | -0.45 | -0.11 | -0.44 | 0.69 |
| Zn | 0.01 | -0.09 | 0.07 | 0.73 | 0.78 | 0.43 | -0.11 | -0.17 | -0.11 | -0.14 | -0.14 |
| Mn | 0.19 | 0.15 | -0.56 | -0.19 | -0.12 | -0.13 | -0.03 | 0.04 | 0.10 | 0.05 | 0.57 |
| Cu | 0.33 | -0.14 | 0.19 | 0.55 | 0.34 | -0.09 | 0.23 | 0.15 | 0.54 | 0.18 | 0.30 |
| B | 0.21 | -0.10 | 0.49 | 0.31 | 0.10 | -0.25 | 0.66 | 0.62 | 0.18 | 0.62 | -0.15 |
| % Sand | -0.55 | 0.11 | -0.62 | -0.19 | 0.13 | 0.51 | -0.68 | -0.82 | -0.81 | -0.83 | -0.32 |
| % Silt | 0.64 | 0.08 | 0.55 | 0.37 | 0.08 | -0.49 | 0.45 | 0.66 | 0.78 | 0.68 | 0.47 |
| % Clay | 0.41 | -0.25 | 0.60 | 0.03 | -0.28 | -0.46 | 0.77 | 0.85 | 0.74 | 0.84 | 0.17 |
| % Organic carbon | 0.50 | -0.17 | 0.60 | 0.62 | 0.38 | -0.39 | 0.67 | 0.76 | 0.66 | 0.78 | 0.15 |

| **Table S9 continued.** | |  |  |  |  |  |  |  |
| --- | --- | --- | --- | --- | --- | --- | --- | --- |
|  | Correlation coefficients | | | | | | | |
| Soil characteristic | Fe | Zn | Mn | Cu | B | % Sand | % Silt | % Clay |
| Moisture |  |  |  |  |  |  |  |  |
| Temp |  |  |  |  |  |  |  |  |
| pH |  |  |  |  |  |  |  |  |
| EC |  |  |  |  |  |  |  |  |
| NON_3_ |  |  |  |  |  |  |  |  |
| P |  |  |  |  |  |  |  |  |
| K |  |  |  |  |  |  |  |  |
| Ca |  |  |  |  |  |  |  |  |
| Mg |  |  |  |  |  |  |  |  |
| S |  |  |  |  |  |  |  |  |
| Na |  |  |  |  |  |  |  |  |
| Fe | 1 |  |  |  |  |  |  |  |
| Zn | -0.05 | 1 |  |  |  |  |  |  |
| Mn | 0.74 | -0.17 | 1 |  |  |  |  |  |
| Cu | 0.18 | 0.30 | 0.15 | 1 |  |  |  |  |
| B | -0.32 | -0.05 | 0.09 | 0.13 | 1 |  |  |  |
| % Sand | 0.24 | 0.25 | -0.10 | -0.38 | -0.51 | 1 |  |  |
| % Silt | -0.15 | -0.08 | 0.08 | 0.46 | 0.31 | -0.91 | 1 |  |
| % Clay | -0.29 | -0.35 | 0.11 | 0.27 | 0.61 | -0.95 | 0.73 | 1 |
| % Organic carbon | -0.24 | 0.19 | 0.11 | 0.60 | 0.65 | -0.80 | 0.78 | 0.73 |

| **Table S10.** Principal Component Analysis (PCA) loadings for soil characteristics across sampling sites, across species. Loadings indicate the contribution of each soil trait to the first three principal components, which together explain 63.5% of the total variation in environmental niche space. Variables with higher absolute loadings contribute more strongly to each axis. | | | |
| --- | --- | --- | --- |
| Loading | PCA axis | | |
|  | PC1 (39.7%) | PC2 (18.2%) | PC3 (5.6%) |
| Moisture | 0.502 | 0.697 | 0.053 |
| Temp | -0.204 | 0.339 | -0.064 |
| pH | 0.754 | -0.494 | 0.254 |
| EC | 0.351 | 0.292 | 0.861 |
| NON_3_ | 0.033 | 0.328 | 0.854 |
| P | -0.505 | -0.227 | 0.370 |
| K | 0.778 | -0.483 | -0.062 |
| Ca | 0.895 | -0.282 | -0.121 |
| Mg | 0.792 | 0.143 | -0.173 |
| S | 0.907 | -0.257 | -0.096 |
| Na | 0.136 | 0.921 | -0.291 |
| Fe | -0.374 | 0.765 | -0.271 |
| Zn | -0.092 | 0.101 | 0.890 |
| Mn | 0.005 | 0.544 | -0.433 |
| Cu | 0.439 | 0.424 | 0.349 |
| B | 0.638 | -0.189 | 0.092 |
| % Sand | -0.955 | -0.117 | 0.200 |
| % Silt | 0.861 | 0.326 | -0.034 |
| % Clay | 0.914 | -0.063 | -0.307 |
| % Organic carbon | 0.908 | 0.150 | 0.288 |

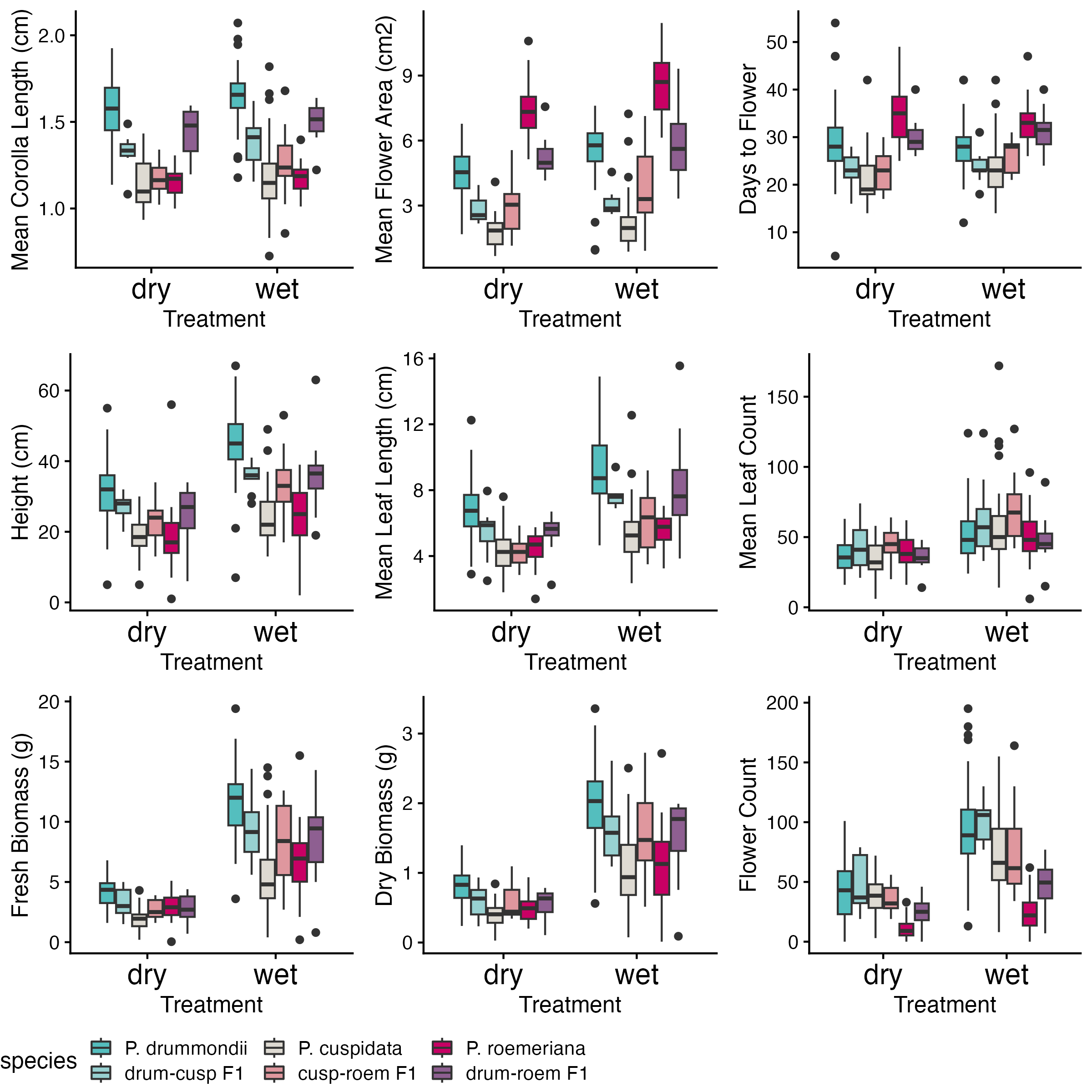

**Figure S2.** Trait distributions for the drydown experiment, separated by treatment and species, including hybrids. Boxplots show medians and interquartile range with potential outlier points.

| **Table S11.** Principal Component Analysis (PCA) loadings for dry-down morphological and phenological traits across the three species. Loadings indicate the contribution of each soil trait to the first three principal components, which together explain 82.8% of the total variation in environmental niche space. Variables with higher absolute loadings contribute more strongly to each axis. | | | |
| --- | --- | --- | --- |
| Loading | PCA axis | | |
|  | PC1 (51.4%) | PC2 (17.9%) | PC3 (13.5%) |
| Mean Corolla Length | 0.753 | 0.050 | -0.451 |
| Mean Flower Area | 0.466 | 0.714 | 0.057 |
| Days to Flower | 0.221 | 0.809 | 0.288 |
| Height | 0.856 | -0.015 | -0.311 |
| Mean Leaf Length | 0.840 | 0.124 | -0.185 |
| Mean Leaf Count | 0.352 | -0.274 | 0.838 |
| Fresh Biomass | 0.942 | -0.025 | 0.215 |
| Dry Biomass | 0.946 | -0.120 | 0.217 |
| Flower Count | 0.688 | -0.590 | -0.048 |

| **Table S12**. Results of Tukey’s HSD test comparing pairwise species differences under dry and wet treatments. Estimates, standard errors (SE), degrees of freedom (df), t-values (*t*), and *P*-values are shown for each trait comparison between species pairs: *Phlox cuspidata* vs *Phlox drummondii* (C-D), *Phlox cuspidata* vs *Phlox roemeriana* (C-R), and *Phlox drummondii* vs *Phlox roemeriana* (D-R). Bolded *P*-values indicate significance (*P* < 0.05). | | | | | | | | | | | |
| --- | --- | --- | --- | --- | --- | --- | --- | --- | --- | --- | --- |
|  | | Dry | | | | | Wet | | | | |
| Trait | Comparison | Estimate | SE | df | *t* | *P* | Estimate | SE | df | *t* | *P* |
| Height | C-D | -8.536 | 3.360 | 34.9 | -2.544 | 0.3479 | -16.174 | 3.400 | 34.5 | -4.761 | 0.17 |
|  | C-R | 4.681 | 3.330 | 92.4 | 1.407 | 0.9594 | 5.120 | 3.260 | 88.8 | 1.569 | 0.9154 |
|  | **D-R** | **13.216** | **3.720** | **33.3** | **3.553** | **0.045** | **21.294** | **3.710** | **32.5** | **5.733** | **0.0001** |
| Leaf Length | C-D | -2.147 | 0.577 | 30.9 | -3.723 | 0.0315 | -3.588 | 0.580 | 29.5 | -6.183 | 0.1 |
|  | C-R | -0.066 | 0.607 | 59 | -0.108 | 1 | -0.228 | 0.596 | 55.5 | -0.383 | 1 |
|  | D-R | 2.081 | 0.630 | 29.7 | 3.32 | 0.848 | 3.360 | 0.630 | 28.6 | 5.337 | 0.5 |
| Leaf Count | C-D | -5.226 | 6.120 | 30.6 | -0.855 | 0.9991 | -0.596 | 6.150 | 29.3 | -0.097 | 1 |
|  | C-R | -1.190 | 6.550 | 56.4 | -0.182 | 1 | 9.120 | 6.400 | 52 | 1.425 | 0.9529 |
|  | D-R | 4.036 | 6.750 | 30.1 | 0.598 | 1 | 9.716 | 6.720 | 28.7 | 1.446 | 0.9435 |
| Fresh Biomass | **C-D** | **-2.275** | **0.625** | **28.4** | **-3.639** | **0.0409** | **-6.115** | **0.626** | **26.5** | **-9.765** | **< 0.0001** |
|  | C-R | -0.911 | 0.697 | 42.9 | -1.307 | 0.9737 | -1.079 | 0.678 | 39 | -1.591 | 0.9019 |
|  | D-R | 1.364 | 0.683 | 28.6 | 1.998 | 0.6913 | **5.036** | **0.679** | **27** | **7.421** | **< 0.0001** |
| Dry Biomass | C-D | -0.423 | 0.120 | 29 | -3.518 | 0.0532 | **-0.963** | **0.120** | **26.3** | **-8.044** | **< 0.0001** |
|  | C-R | -0.094 | 0.135 | 45.1 | -0.698 | 0.9999 | -0.055 | 0.129 | 38.9 | -0.423 | 1 |
|  | D-R | 0.329 | 0.132 | 29.9 | 2.493 | 0.3813 | **0.908** | **0.130** | **26.8** | **7.003** | **< 0.0001** |
| Flower Count | C-D | -5.381 | 6.140 | 29.9 | -0.877 | 0.9989 | **-22.959** | **6.080** | **26.6** | **-3.775** | **0.0315** |
|  | **C-R** | **25.541** | **7.030** | **45.7** | **3.633** | **0.0304** | **47.414** | **6.760** | **38.8** | **7.017** | **< 0.0001** |
|  | **D-R** | **30.922** | **6.740** | **32.5** | **4.589** | **0.0031** | **70.373** | **6.610** | **28.6** | **10.643** | **< 0.0001** |
| Days to Flower | **C-D** | **-6.405** | **1.510** | **28.4** | **-4.252** | **0.0093** | -3.605 | 1.510 | 25.7 | -2.395 | 0.443 |
|  | **C-R** | **-12.846** | **1.720** | **45.3** | **-7.475** | **< 0.0001** | **-9.065** | **1.670** | **40.2** | **-5.428** | **0.0002** |
|  | **D-R** | **-6.440** | **1.670** | **31.6** | **-3.858** | **0.0222** | -5.461 | 1.650 | 28.4 | -3.319 | 0.0836 |
| Corolla Length | **C-D** | **-0.413** | **0.047** | **27.7** | **-8.733** | **< 0.0001** | **-0.460** | **0.046** | **23.2** | **-9.97** | **< 0.0001** |
|  | C-R | -0.008 | 0.062 | 64.8 | -0.121 | 1 | 0.004 | 0.052 | 39.6 | 0.075 | 1 |
|  | **D-R** | **0.405** | **0.062** | **47** | **6.574** | **< 0.0001** | **0.464** | **0.052** | **26.7** | **8.976** | **< 0.0001** |
| Flower Area | **C-D** | **-1.824** | **0.451** | **31.3** | **-4.041** | **0.0141** | **-2.643** | **0.449** | **27.6** | **-5.884** | **0.0001** |
|  | **C-R** | **-4.882** | **0.487** | **56.8** | **-10.022** | **< 0.0001** | **-5.786** | **0.472** | **49** | **-12.265** | **< 0.0001** |
|  | **D-R** | **-3.058** | **0.502** | **31.4** | **-6.089** | **0.0001** | **-3.143** | **0.490** | **27** | **-6.49** | **< 0.0001** |

| **Table S12 continued.** | | | | | | |
| --- | --- | --- | --- | --- | --- | --- |
|  | | Within Species | | | | |
| Trait | Comparison | Estimate | SE | df | t | *P* |
| Height | **Dry C-Wet C** | **-5.967** | **1.570** | **294.4** | **-3.789** | **0.0098** |
|  | **Dry D-Wet D** | **-13.605** | **1.390** | **308.4** | **-9.798** | **<.0001** |
|  | Dry R-Wet R | -5.527 | 1.860 | 299.2 | -2.971 | 0.1224 |
| Leaf Length | Dry C-Wet C | -0.975 | 0.327 | 307.2 | -2.982 | 0.119 |
|  | **Dry D-Wet D** | **-2.417** | **0.281** | **308.9** | **-8.603** | **<.0001** |
|  | Dry R-Wet R | -1.138 | 0.385 | 312.4 | -2.96 | 0.1258 |
| Leaf Count | **Dry C-Wet C** | **-21.788** | **3.580** | **294.5** | **-6.087** | **<.0001** |
|  | **Dry D-Wet D** | **-17.158** | **3.120** | **307.1** | **-5.502** | **<.0001** |
|  | Dry R-Wet R | -11.479 | 4.290 | 299 | -2.673 | 0.2449 |
| Fresh Biomass | **Dry C-Wet C** | **-3.607** | **0.463** | **306.3** | **-7.788** | **<.0001** |
|  | **Dry D-Wet D** | **-7.446** | **0.400** | **314.8** | **-18.62** | **<.0001** |
|  | **Dry R-Wet R** | **-3.775** | **0.545** | **313** | **-6.923** | **<.0001** |
| Dry Biomass | **Dry C-Wet C** | **-0.638** | **0.088** | **301.9** | **-7.228** | **<.0001** |
|  | **Dry D-Wet D** | **-1.178** | **0.076** | **310.5** | **-15.435** | **<.0001** |
|  | **Dry R-Wet R** | **-0.599** | **0.105** | **307** | **-5.711** | **<.0001** |
| Flower Count | **Dry C-Wet C** | **-35.019** | **5.270** | **307.6** | **-6.643** | **<.0001** |
|  | **Dry D-Wet D** | **-52.598** | **4.550** | **312.1** | **-11.569** | **<.0001** |
|  | Dry R-Wet R | -13.146 | 6.250 | 314.6 | -2.105 | 0.62 |
| Days to Flower | Dry C-Wet C | -2.392 | 1.190 | 299.5 | -2.006 | 0.6891 |
|  | Dry D-Wet D | 0.409 | 1.030 | 301.9 | 0.399 | 1 |
|  | Dry R-Wet R | 1.389 | 1.460 | 303.2 | 0.952 | 0.9985 |
| Corolla Length | Dry C-Wet C | -0.044 | 0.035 | 248.3 | -1.248 | 0.9843 |
|  | Dry D-Wet D | -0.091 | 0.031 | 259.9 | -2.971 | 0.1232 |
|  | Dry R-Wet R | -0.032 | 0.054 | 258.2 | -0.599 | 1 |
| Flower Area | Dry C-Wet C | -0.309 | 0.266 | 261.8 | -1.164 | 0.9911 |
|  | **Dry D-Wet D** | **-1.128** | **0.228** | **266.6** | **-4.959** | **0.0001** |
|  | **Dry R-Wet R** | **-1.213** | **0.329** | **263.2** | **-3.691** | **0.0141** |

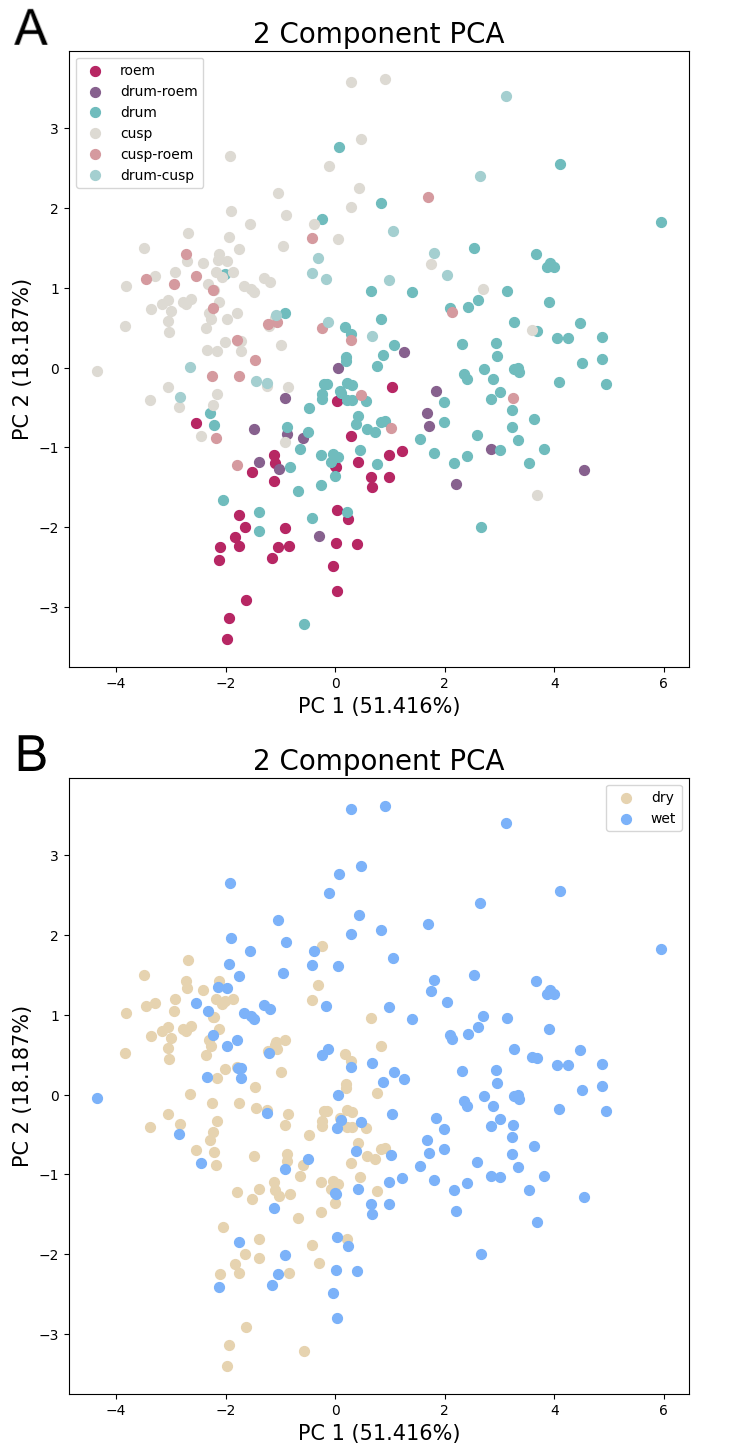

**Figure S3.** (A) PCA plot of Phlox *drummondii*, *cuspidata*, and *roemeriana* individuals, as well as their hybrids, based on measurements taken during the drydown experiment, colored by species. Dimensionality reduction was performed on mean corolla length, mean flower area, height, mean leaf length, leaf count, biomass (fresh and dry), days to flower, and flower count. (B) The same PCA plot, colored by drydown treatment.
